# Using deep learning predictions of inter-residue distances for model validation

**DOI:** 10.1101/2022.08.25.505292

**Authors:** Filomeno Sánchez Rodríguez, Grzegorz Chojnowski, Ronan M. Keegan, Daniel J. Rigden

## Abstract

Determination of protein structures typically entails building a model that satisfies the collected experimental observations and its deposition in the Protein Data Bank (PDB). Experimental limitations can lead to unavoidable uncertainties during the process of model building, which result in the introduction of errors into the deposited model. Many metrics are available for model validation, but most are limited to the consideration of the physico-chemical aspects of the model or its match to the map. The latest advances in the field of deep learning have enabled the increasingly accurate prediction of inter-residue distances, an advance which has played a pivotal role in the recent improvements observed in the field of protein *ab initio* modelling. Here we present new validation methods based on the use of these precise inter-residue distance predictions, which are compared with the distances observed in the protein model. Sequence register errors are particularly clearly detected, and the register shifts required for their correction can be reliably determined. The method is available in the package ConKit (www.conkit.org).

## 1. Introduction

Structural determination of proteins may be carried out using a range of different techniques, of which macromolecular X-ray crystallography (MX) and cryogenic electron microscopy (cryoEM) are currently the most popular. These experiments typically culminate in the creation of a model that satisfies the experimental observations collected, and which is later deposited in the Protein Data Bank (PDB) (Berman et al., 2000). However, as in all experiments, these observations will have unavoidable uncertainties caused by experimental limitations, which can result in the introduction of errors into the final model.

Such errors were particularly common during the early stages of X-ray crystallography, when technical advances allowed for an increasing number of protein folds to be experimentally determined and deposited in the PDB. It was also during this time that some deposited structures were first found to contain major errors, highlighting the need for model validation tools (Hooft et al., 1996; Kleywegt & Jones, 1995; MacArthur et al., 1994). Several computational methods and systems were developed to address the issue. Among them were PROCHECK (Laskowski et al., 1993) and WHATIF (Vriend, 1990), each majoring on geometric and stereochemical properties and generating residue-by-residue reports to inform the user of potential errors in the model. However, it was noted that stereochemical analyses could be insufficient for unambiguous identification of errors leading to the introduction of methods such as VERIFY_3D (Lüthy et al., 1992) based on statistics of favoured amino-acid environments, ProSA (Sippl, 1993) based on the combination of a Cβ-Cβ (or Cα-Cα) potential and solvent exposure statistics, and ERRAT (Colovos & Yeates, 1993) which is based on the statistics of nonbonded interactions between C, N and O atoms. Further notable contributions include COOT (Emsley & Cowtan, 2004), which provided interactive model building tools coupled with a series of residue-by-residue model validation metrics based both on the geometric properties of the model and its match to the map. MolProbity (Davis et al., 2007) was later released and provided validation reports based on the analysis of all-atoms contacts together with other geometric and dihedral-angle analyses.

With the recent technical improvements in image processing and electron detectors a rapid increase in the number of molecules deposited in the PDB which were solved using cryoEM has been observed in recent years (Chiu et al., 2021). Compared with X-ray crystallography, these models are often built using lower resolution data, with maps that have varying levels of local resolution at different parts of the model, all of which can hinder model building and make cryoEM models more susceptible to errors. The discovery of modelling errors among cryoEM structures recently deposited in the PDB (Chojnowski et al., 2022; Croll et al., 2021; Weiss et al., 2016), has highlighted the need for new tools for model validation (Afonine et al., 2018; Lawson & Chiu, 2018). This has led to the creation of sophisticated new tools for model validation, such as checkMySequence (Chojnowski, 2022), which uses the latest advances in machine learning for the detection of out-of-register sequence errors. New metrics for the assessment of model quality and its fit to the map have also been introduced in recent years, such as SMOC (Joseph et al., 2016), a segment-based Manders’ overlap coefficient between the model and the map, FSC-Q (Ramírez-Aportela et al., 2021) a model quality validation score based on the local Fourier shell correlation between the model and the map, and the map Q-Score (Pintilie et al., 2020), which measures atomic resolvability. Creation of these metrics has resulted in the development of new user interfaces to integrate these different metrics in order to facilitate their interpretability and provide an all-in-one package. This is the case of the CCP-EM validation task (Joseph et al., 2022), which combines several of these new metrics and tools into a graphical user interface. Further developments in interpretability of validation metrics came with the release of Iris (Rochira & Agirre, 2021), a tool that combines different validation metrics calculated on a residue-by-residue basis.

Recent developments in the field of evolutionary covariance and machine learning have enabled the precise prediction of residue-residue contacts and increasingly accurate inter-residue distance predictions (Ruiz-Serra et al., 2021). Access to this accurate covariance information has played an essential role in the latest advances observed in the field of protein structural bioinformatics, particularly the improvement of protein *ab initio* modelling, with the most notable examples being AlphaFold 2 (Jumper et al., 2021) and RoseTTAFold (Baek et al., 2021).

Here we present new validation methods based on the availability of accurate inter-residue distance predictions. Potential errors are recognised as residues and regions for which contacts and inter-residue distances observed in the model differ significantly from those predicted by Deep Learning-based methods. A series of metrics relating to the consistency of observed and predicted contacts and distances are fed into a support-vector machine classifier that was trained to detect model errors using historical data from the EM Validation Challenges (Lawson et al., 2021; Lawson & Chiu, 2018). Further detection of possible register errors specifically is done by performing an alignment of the predicted contact map and the map inferred from the contacts observed in the model. Regions of the model where the maximum contact overlap is achieved through a sequence register different to that observed in the model are flagged and the optimal sequence register can then be used to fix the register error. Results suggest that the detection of model errors and the correction of sequence register errors is possible through the use of the trained classifier in conjunction with the contact map alignment, as revealed by analysing a set of structures deposited in the PDB. The approach, implemented in ConKit (Simkovic et al., 2017) through the command line option “conkit-validate”, thus provides a new tool for protein structure validation that is orthogonal to existing methods.

## 2. Materials and methods

### 2.1 Creation of a training dataset of misregistered residues extracted from the EM modelling challenges

Structures submitted to the EM modelling challenges that took place in 2016, 2019 and 2021 (Lawson et al., 2021; Lawson & Chiu, 2018) were analysed in order to create a database containing modelling errors, annotated according to whether the cause was or was not an incorrect sequence register. First, structures where more than half of residues scored a sequence-dependent local-global alignment (LGA) (Zemla, 2003) above 8Å between the target and the experimentally determined structure were discarded. For the remaining structures, LGA values were smoothed using a three residue-window rolling average. Model regions where at least 3 consecutive residues scored a smoothed LGA value of 3Å or higher were then visually inspected searching for register errors. These errors were defined as ranges of residues where despite having a smoothed LGA above 3Å, the main chain had been modelled correctly when compared with the ground-truth solution. To reduce redundancy, register errors found within the same sets of residues across different models submitted for the same target were removed, except for the error affecting the largest number of residues, which was selected as the representative error. All residues found among the resulting set of register errors were then labelled with the positive class (modelled incorrectly) and taken into a database of register errors. The remaining residues found in the models from which these register errors were taken were also added into the database, but they were labelled instead with the negative class (modelled correctly) if the smoothed LGA was below 3Å, otherwise they were considered part of a modelling error and the positive class was assigned (modelled incorrectly). This resulted in the creation of a dataset consisting of 8620 residues, of which 6192 were labelled with the negative class and 2428 with the positive class. Residues labelled with the positive class were extracted out of 76 sequence register errors (2278 residues) and 12 other modelling errors (150 residues).

### 2.2 Prediction of inter-residue distances using AlphaFold 2

Predictions of inter-residue contacts and distances were obtained for the models being validated using AlphaFold 2 (Jumper et al., 2021). The original CASP14 model preset was used and the database search was set to full mode. All other parameters were left with their default values. Predictions were carried out on a computing grid where each node is equipped with a twin 16-core Intel Xeon Gold 5218 running at 2.3GHz, 187GB of memory and four NVIDIA Tesla V100 chips with 16GB of video memory each.

For each case, five models were produced and the inter-residue distance predictions for the model with the highest predicted Local Distance Difference Test (pLDDT) (Mariani et al., 2013) were taken. For each residue pair in the structure, this prediction contains the predicted probability that these residues are within a series of distance bins. These distances were processed for all residue pairs so that the midpoint value of the distance bin with the highest probability was considered the predicted distance, and the probability associated with this bin the confidence score. Contact predictions were derived from the distance prediction by adding together all the probabilities observed across the distance bins up to 8Å. The top L/2 contacts scoring the highest probability values were then taken to form the final predicted contact maps, where L denotes the sequence length.

### 2.3 New covariance-based metrics for model validation and feature engineering

A set of new metrics were developed with the aim of comparing the inter-residue contacts and distances observed in a model and those predicted for the protein of interest. Rather than a global comparison of the similarity of two contact maps or distograms, these metrics were designed for local model validation, hence they are calculated on a residue-by-residue basis.

First, a weighted root mean square deviation (wRMSD) of the predicted and the observed inter-residue distances was calculated for each residue of the model as follows:

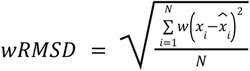

Where N represents the number of residues in the model, x the observed distance for a pair of residues, 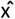 the predicted distance for a pair of residues, and w the confidence of the predicted inter-residue distance (a value between 0 and 1).

A series of metrics based on the analysis of the inter-residue contacts on a residue-by-residue basis were also defined as follows:

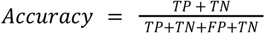

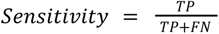

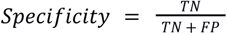

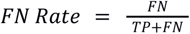

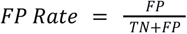

Where TP represents true positives, FP false positives, TN true negatives and FN false negatives. Additionally, the raw count of FN and FP was also used as two additional features. Two residues are considered in contact with each other when their Cβ atoms are within 8Å of each other. For the calculation of these metrics, the top L/2 contacts scoring the highest confidence values were used - where L denotes the number of residues in the protein sequence. Additionally, the raw count values of FN and FP were also used as a metric.

For all the proposed metrics, smoothed and unsmoothed versions were calculated. Whereas in the un-smoothed version the values observed across the residues of the model were kept intact, for the smoothed version these values were smoothed using a convolution approach. In this approach, a five point unweighted filter was used to convolve the raw data, making this transformation equivalent to using a moving five residue window averaging technique, with the added benefit of not losing data on the edges of the model where there’s not enough information to calculate a window average.

Additionally, for all the proposed metrics a Z-Score derived metric was computed. This was calculated taking the value observed for each residue of the model, and using the values observed for the residues within a range of 10Å as the full sample.

This resulted in the creation of 24 metrics: one distance prediction-based metric, 7 contact prediction-based metrics, their smoothed and un-smoothed versions, and the additional Z-Score version. To ensure minimal autocorrelation, Pearson’s correlation coefficients were examined for all the possible pairs of these metrics using the values observed across the EM modelling challenge dataset described in Section 2.1 (Figure 1). Where a pair or a group of metrics shared an absolute correlation value of 0.4 or higher, only one representative metric was taken and the others discarded. The representative metric was selected based on initial results obtained using a linear discriminant analysis using the residues from the EM challenge dataset and setting the class label as the target. For a given set of correlated metrics, the feature with the highest coefficient observed in this analysis was selected as the representative. This resulted in the creation of a final set of 7 metrics: Accuracy, FP Rate, Smooth Sensitivity, Smooth wRMSD, Z-Score Accuracy, Z-Score Sensitivity, and Z-Score wRMSD.

**Figure 1.**
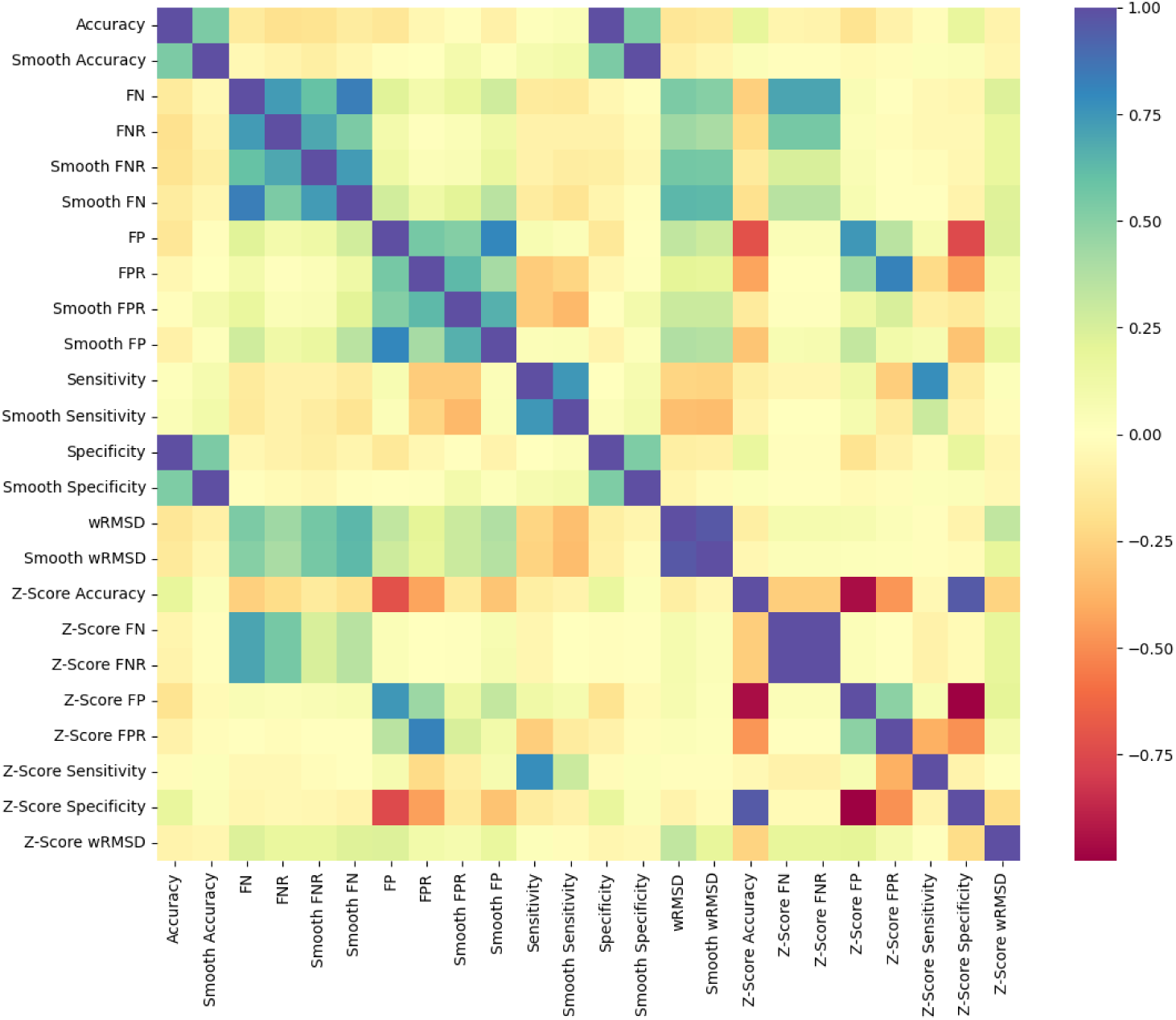
Correlation matrix for the proposed covariance-based metrics for model validation. Colour at each cell corresponds with the Pearson’s correlation coefficient observed between each pair of metrics. The scale goes from red for negative correlation to deep blue for positive correlation.

### 2.4 Machine learning training and hyperparameter tuning

Residues found in the dataset created by extracting residues from the EM modelling challenge submissions were used to train several machine learning algorithms for the detection of register errors. For each observation in this dataset, covariance metrics described in the previous Section 2.3 were calculated. Additional features describing the residue’s local environment were also included into the observations, specifically the residue solvent accessibility (ACC) and the secondary structure element where the residue was located (Helix, Beta Sheet or Coil). Residues found to be part of register and other modelling errors were labelled with the positive target class otherwise with the negative class. To create the training and test set, residues labelled with the positive class were randomly split using a 80:20 ratio. In order to ensure balanced datasets, the same number of residues labelled with the negative class were randomly selected and added into each set. This resulted in the creation of a training set consisting of 3884 observations and a test set formed by 972 observations, both of them having a balanced number of observations in each class. The data was then scaled to unit standard deviation and a zero-mean using a standard scaler, fitted only using the data seen in the training set to prevent data leakage. Optimal training hyperparameters for each classifier were found by performing a random search of 200 iterations using the mean accuracy as the scoring function. All the algorithm implementations were done with scikit-learn version 0.24.2 (Pedregosa et al., 2011).

### 2.5 Contact map alignment-based sequence re-assignment

The alignment between predicted contact maps and the contact maps observed in the models of interest was calculated using *map_align* (Ovchinnikov et al., 2017). This tool creates an alignment between two input contact maps so that the contact maximum overlap (CMO) is achieved (Andonov et al., 2011), by introducing and extending gaps as necessary. The CMO is defined as the number of matching contacts between the two input contact maps, when optimal alignment is achieved. If a misalignment between the input contact maps is found for a set of residues, the sequence register used to achieve the CMO between predicted and observed contacts is proposed as a fix for the possible register error.

### 2.6 Creation of a benchmark dataset consisting of PDB-deposited structures solved with cryoEM

Protein structures determined using cryo-EM at 5Å resolution or better, with or without a nucleic acid component were selected from the Protein Data Bank (PDB) (Berman et al., 2000) as of 10 November 2021. For practical reasons out of 7241 available structures a subset of 5744 was selected with size of corresponding compressed EM maps not exceeding 200MB. Next, all the structures were automatically analysed using the checkMySequence validation tool (Chojnowski, 2022) that identified 419 chains with tentative sequence assignment issues in 246 structures that were used for further analysis.

Using the reference sequence deposited in the PDB, distance predictions were obtained for each individual chain. For 55 chains this was not possible due to hardware limitations, particularly the system running out of memory before the predictions could be complete. Ultimately, our analysis could be applied to 364 protein chains found in structures deposited in the PDB where possible register errors were reported by checkMySequence. To ensure a match between the residue numbering observed in the deposited models and the numbering in the reference sequence used to obtain the inter-residue distance predictions, CROPS (https://github.com/rigdenlab/crops) was used to renumber the models based on the reference. For the 149 cases where this was not possible due to major inconsistencies between the protein sequence and residue numbering in the deposited model, a manual inspection was carried out.

## 3. Results and Discussion

### 3.1 Machine learning detects register errors using covariance-based metrics

In order to create a classifier able to detect register and other modelling errors using the newly developed covariance-based metrics, three different types of classifiers available at scikit-learn (Pedregosa et al., 2011) were selected: Support Vector Machines using a linear kernel (SVM), Random Forest (RF) and K-nearest Neighbours (KNN). To create the training and test set, residues from the models submitted to the EM modelling challenges were extracted, and a train-test split was created as described in Sections 2.1 and 2.4. Optimal training hyperparameters found in a random search were then used to train each of the classifiers using the observations in the training set, and prediction of the test set was then attempted. Analysis of the results obtained after the prediction of the test samples with each of the classifiers revealed an overall good performance by all three classifiers, which were able to provide accurate predictions for the most part of residues present in the test set (Figure 2). Further analysis was done by plotting the receiver operating characteristic (ROC) curves of each classifier (Figure 3), which showed that the three classifiers performed well at different confidence score threshold cutoff values.

**Figure 2.**
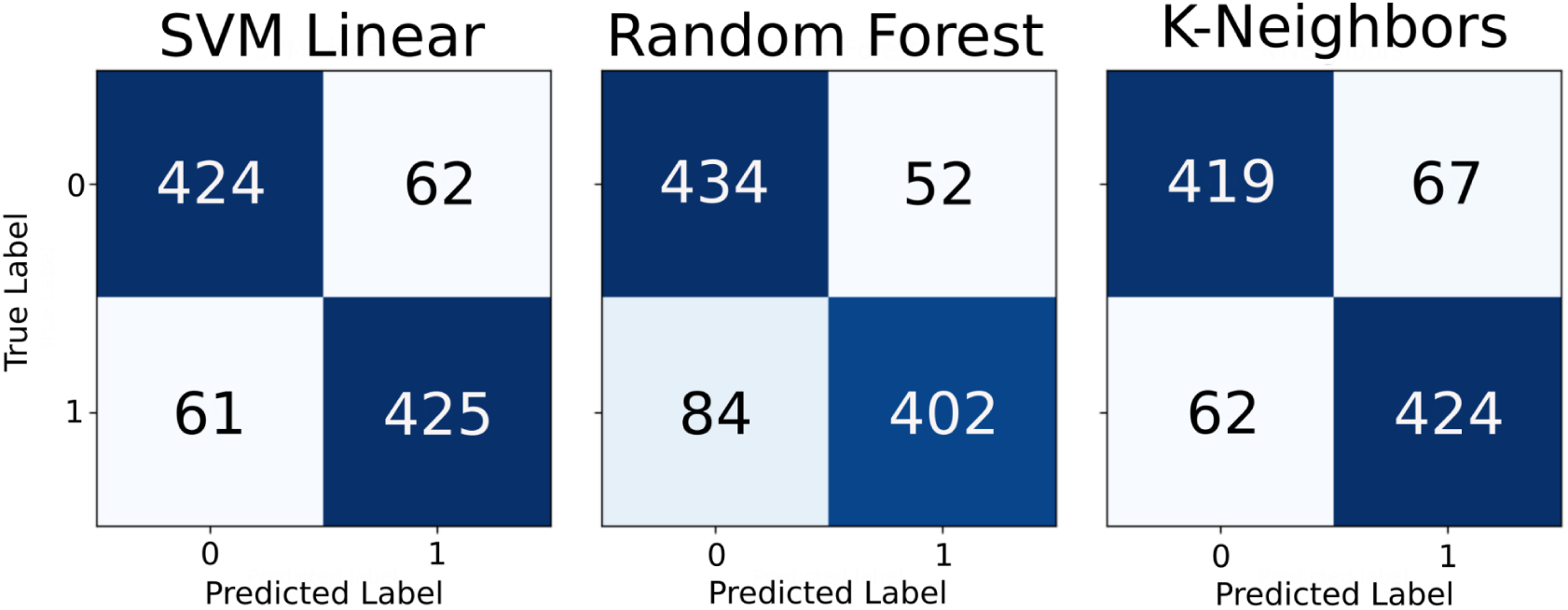
Confusion matrices obtained after predicting the data points at the test set with each of the three classifiers being assessed: Support Vector Machines, Random Forest and K-nearest Neighbours. Residues present in a register error are labelled as class 1, residues modelled correctly as class 0. Numbers inside each cell correspond with the number of data points in each category, which is proportional to the colour intensity at each cell.

**Figure 3.**
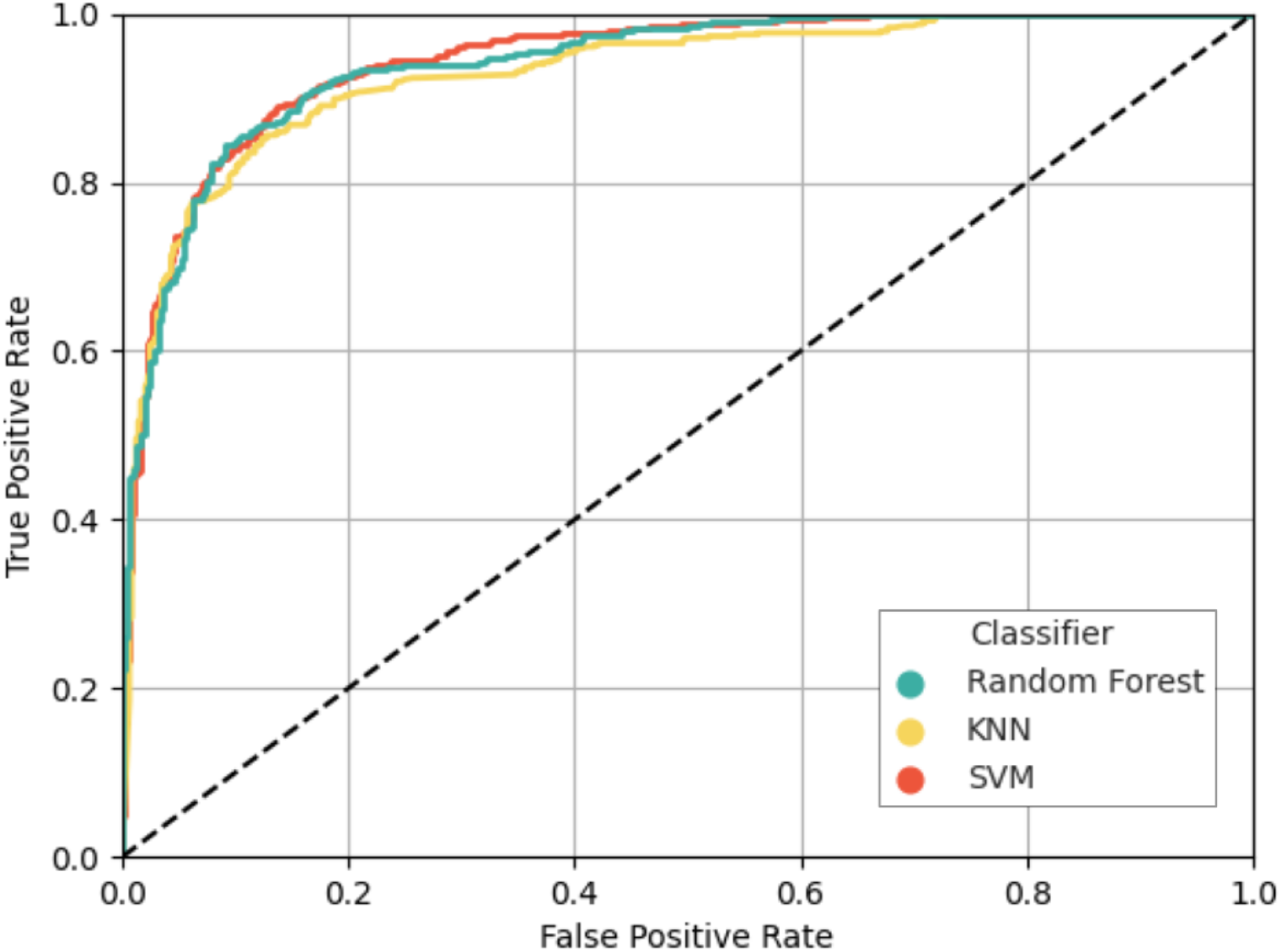
Receiver operating characteristic (ROC) curve depiction of the results obtained on the test set using each of the three classifiers being assessed: Support Vector Machines (orange), Random Forest (yellow) and K-nearest Neighbours (turquoise). Horizontal and vertical axes correspond with the two operating characteristics, the False Positive rate and True Positive Rate respectively, measured at different confidence threshold values. A dashed diagonal line represents the performance of a no-skill classifier where predictions are made at random.

The performance of the trained classifiers on the hold-out test set samples was further analysed based on several metrics (Table 1). Interestingly, only some minor differences were observed across the different classifiers, which had a similar overall good performance at predicting whether the residues in the test set were part of model errors or not. Despite achieving the highest precision (0.885), the RF classifier was not the classifier of choice to be integrated into ConKit, the SVM was selected instead after having scored the highest Accuracy, area under the ROC curve (AUC), recall and F1-Score.

**Table 1.**
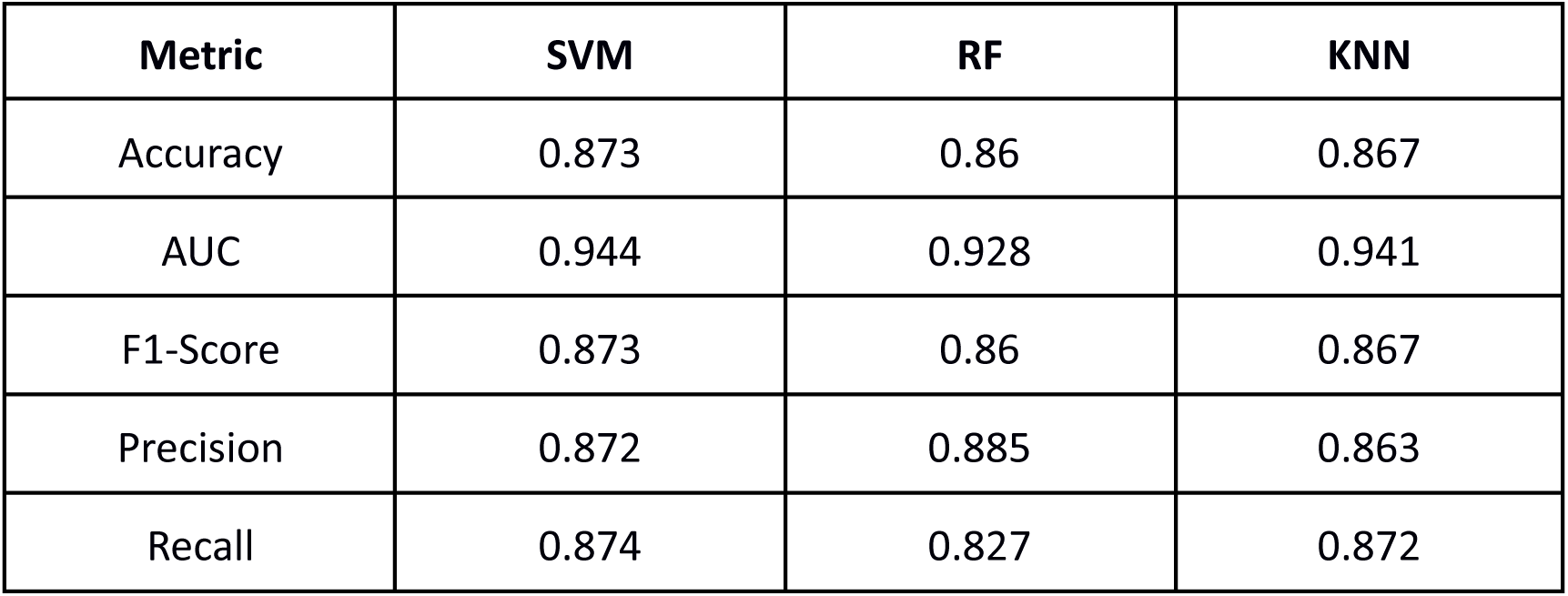
Accuracy, area under the ROC curve (AUC), F1-Score, precision and recall achieved with the hold-out test set by Support Vector Machines (SVM), Random Forest (RF) and K-nearest Neighbours (KNN) classifiers.

| **Metric** | **SVM** | **RF** | **KNN** |
| --- | --- | --- | --- |
| Accuracy | 0.873 | 0.86 | 0.867 |
| AUC | 0.944 | 0.928 | 0.941 |
| F1-Score | 0.873 | 0.86 | 0.867 |
| Precision | 0.872 | 0.885 | 0.863 |
| Recall | 0.874 | 0.827 | 0.872 |

In order to assess the importance of each of the features in the models being compared, a feature permutation analysis was carried out. In this analysis, the values of each feature were randomly shuffled across the samples in the hold-out test set. Prediction was then attempted with the trained classifiers, and the decrease in accuracy with respect to the baseline (accuracy of the trained classifier in the test set without shuffling) was recorded. Each feature permutation was repeated 50 times at random for results consistency. Analysis of the results obtained after these permutations (Figure 4) revealed a strong decrease in accuracy after the shuffling of the wRMSD values, an indication that all three models depend on wRMSD the most for the accurate prediction of whether a given residue is part of a model error or not. This was followed by the sensitivity, which was the second most important feature across the three classifiers. No major differences were observed for the rest of features across the different classifiers, with the exception of the permutation of the Accuracy and Z-Score Accuracy features, which showed a decrease in the performance only for the SVM.

**Figure 4.**
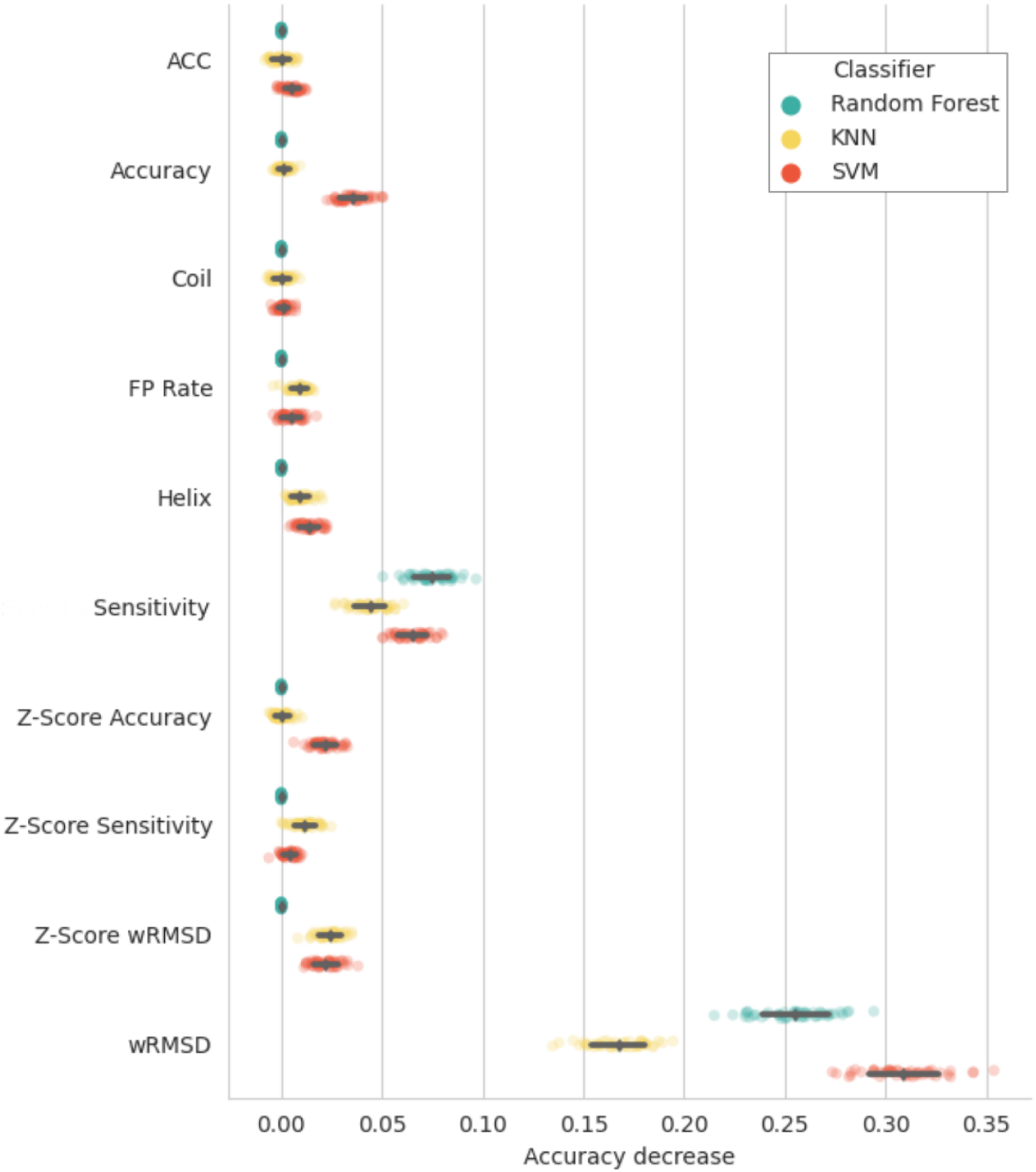
Depiction of the results of the feature permutation analysis for the three classifiers being assessed: Support Vector Machines (orange), Random Forest (yellow) and K-nearest Neighbours (turquoise). Each point represents the accuracy decrease observed after each independent feature permutation. Black diamonds depict the mean accuracy decrease observed for each feature, and the black bars the standard deviation.

The trained SVM classifier outputs a predicted likelihood that a given residue is within a model error (as defined in Section 2.1) on a residue-by-residue basis, meaning that it performs this prediction without any knowledge of the context of the residue of interest. In particular, it does not have any knowledge of the scores predicted for the neighbouring residues. Nevertheless, both sequence register and other kinds of modelling errors are expected to span several consecutive residues within the model being validated. To exploit this expectation, values ranging from 1 to 20 were tested as threshold values of the number of consecutive residues predicted to be within an error. Different thresholds for the score required to predict a residue as an error were tested as well, with the following values being used: 50, 60, 70, 80 and 90%. All the combinations of these values were tested using the models in the EM modelling challenge dataset, and the precision-recall values were recorded (Figure 5). This revealed a negative correlation between the number of consecutive residues required to flag a possible model error and the recall of these errors. Interestingly, higher prediction and recall values were achieved as the score threshold increased, achieving a peak using a threshold of 90%. For instance, a threshold of 3 consecutive residues with a predicted score of at least 50% would achieve a precision of 0.58 and a recall of 0.9, while setting a threshold of 20 consecutive residues with a score of 50% or above would achieve precision and recall scores of 0.81 and 0.3 respectively. Depending on the use-case, users may choose to maximise precision over recall or vice-versa, which is why both thresholds can be tuned through the command line in conkit-validate. Default values were set to a threshold of 6 consecutive residues with a score of 90% or higher to flag possible register errors, which were observed to achieve a precision of 0.92 and a recall of 0.66 on this test.

**Figure 5.**
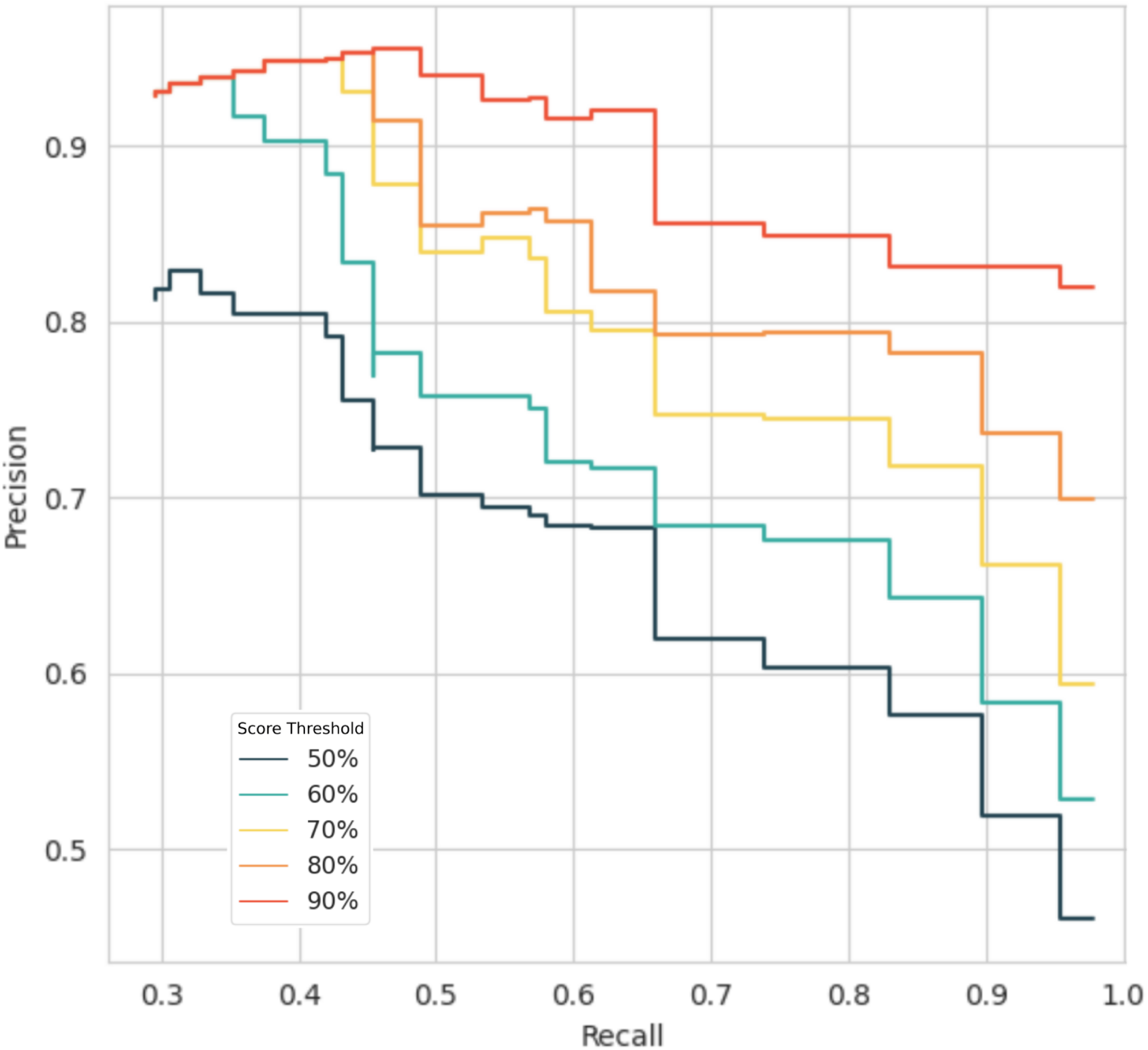
Precision-Recall curves obtained using different threshold values for the number of consecutive residues predicted as errors required to flag a possible register error in a model, and the score required to classify a residue as an error. Curves coloured from dark blue to dark orange correspond with predicted score threshold values of 50, 60, 70, 80 and 90% respectively. For each curve, precision and recall values were recorded after setting different thresholds on the number of consecutive errors required to flag an error: values start at 20 on the left and decrease down to a single residue on the right. Actual values are available at Supplementary Table 1.

### 3.2 Contact map alignment can be used for successful sequence re-assignment of register errors

In order to assess the performance of the contact map alignment as a method to re-assign the correct sequence to register errors, all the models submitted to the EM modelling challenges that form part of the training dataset described in Section 2.1 were tested as follows. The contact maximum overlap (CMO) between the observed contacts in the submitted model and the predicted contact maps was calculated as described in Section 2.5. If for a given range of residues the CMO was achieved using a sequence register different to the one observed in the model being validated (i.e. there is a misalignment between contact maps), then these residues were predicted to be part of a register error and the optimal sequence register was suggested as a fix. Encouragingly, out of all the 88 errors in this dataset, 71 errors were detected using this contact map alignment approach (Figure 6). For all these detected errors, the CMO between the predicted and the observed contacts was achieved using the correct sequence register, which was suggested as a fix, meaning that 87% of the errors analysed could have been fixed if this method had been available for the original authors of the models. Only 17 errors could not be detected using this method. From this, only 5 errors corresponded with register errors, all of them had fewer than 10 residues. The other 12 errors were non-sequence register related modelling errors, for which unsurprisingly, this approach was found to be unsuitable as an alternative sequence register cannot be found to achieve the CMO. Furthermore, only a small number of false positives was observed-instances where the contact map alignment would suggest a different sequence register for the model despite there being no error. Only 6 such cases were found, all of them consisting of misalignments of less than 10 residues.

**Figure 6.**
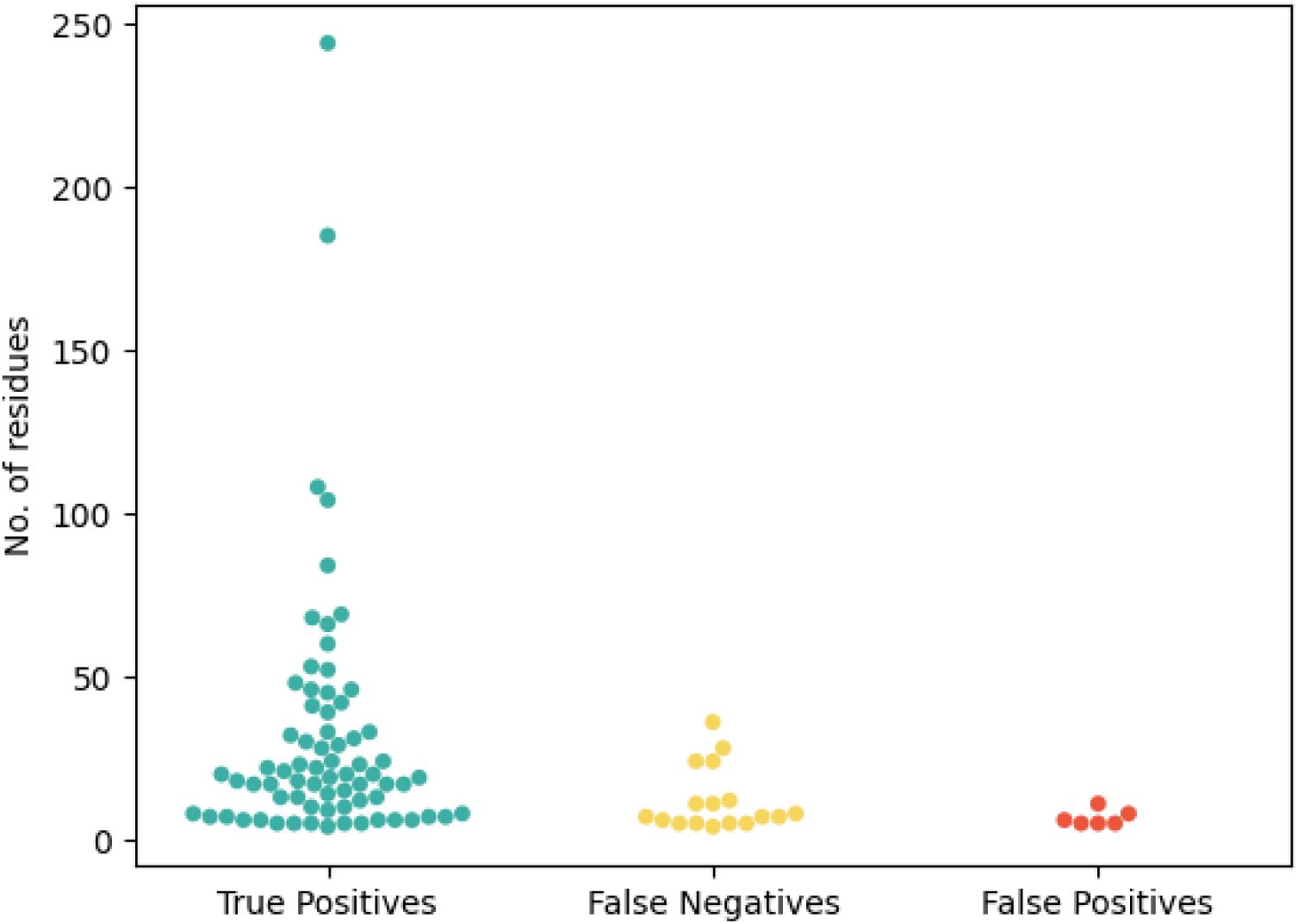
Performance of the sequence re-assignment of register errors in the EM modelling challenges using contact map alignment. The vertical axis shows the number of residues present in the register errors, which are represented with coloured points. True Positives (turquoise) are register errors where the correct sequence register was revealed after the contact map alignment. False negatives (yellow) are register errors where no contact map misalignment was detected at all. False positives (orange) are regions of the models where a contact map misalignment was detected despite there being no register error.

To further characterise those errors that could not be detected using this contact map alignment approach, and those regions of the submitted models that were part of a contact map misalignment despite there being no register error, an analysis of the distribution of residues across the different secondary structure elements was carried out. For each category (True Positives, False Negatives and False Positives), the number of residues found in each secondary structure element was recorded (Figure 7). Initial analysis of this distribution revealed that most register errors are located within beta-sheets, followed by coils and lastly alpha-helices. Interestingly, no difference in the distribution of residues across the three secondary structure elements was evident, all having a high count of TPs, and a significantly smaller count of FN and FP.

**Figure 7.**
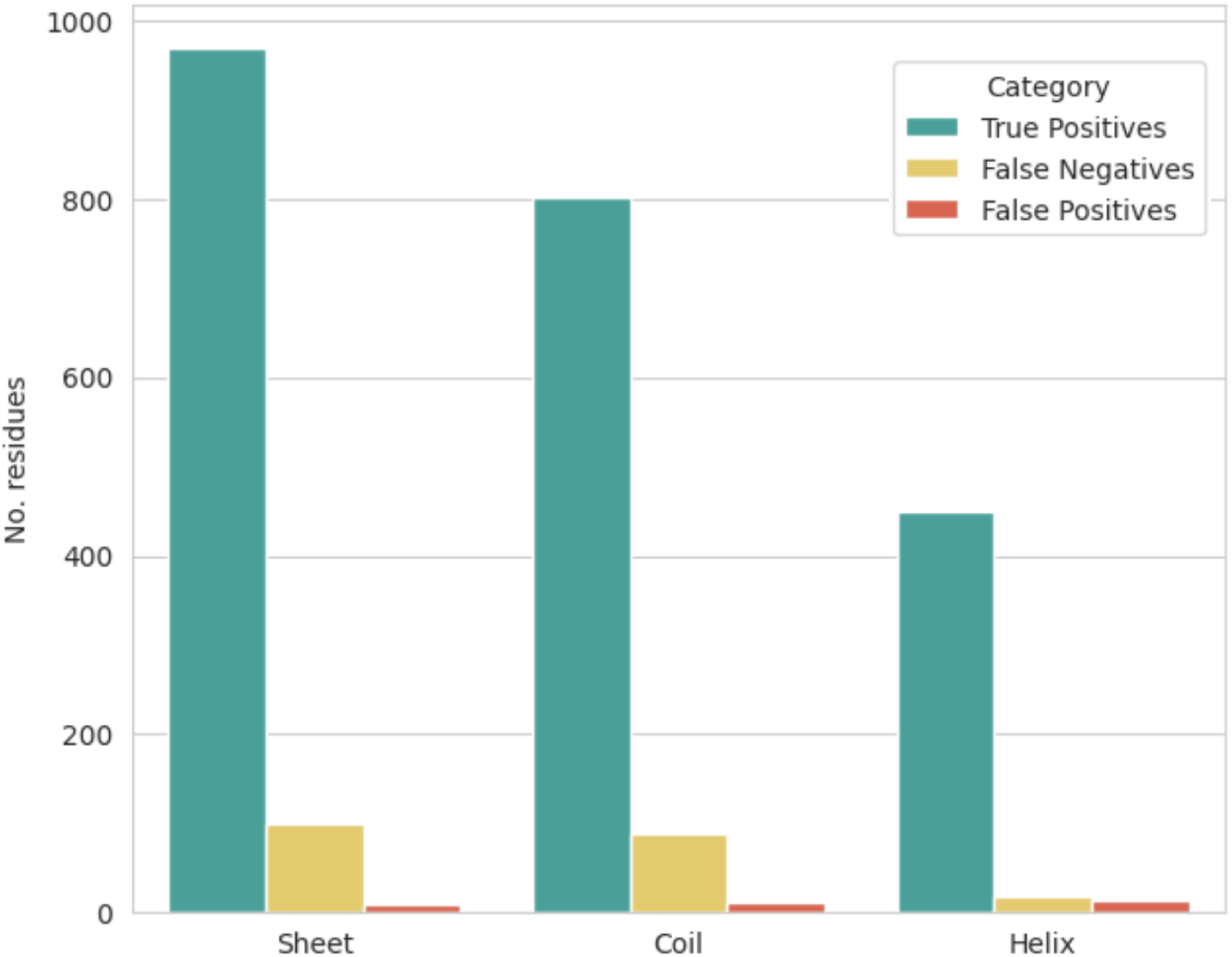
Secondary structure context of residues found in register errors at models submitted to the EM modelling challenges. Vertical axis depicts the number of residues in each category. Residues are grouped along the horizontal axis depending on the secondary structure element. Turquoise bars represent true positives (register errors where the correct sequence register was revealed after the contact map alignment), yellow bars false negatives (register errors where no contact map misalignment was detected at all) and orange bars false positives (regions of the models where a contact map misalignment was detected despite there being no register error).

### 3.3 Contact map alignment and SVM classifier are complementary methods for the detection of modelling errors

In order to assess whether the two proposed methods for model validation complement each other, an analysis of the residues found in register errors that could only be detected using one approach or the other was carried out. In order to do this, the 2428 residues found within register and other modelling errors in the EM modelling challenges, were selected. Out of these, 1943 residues had to be discarded as they were part of the dataset used to train the classifier. This left 486 residues, for which classification was attempted using the SVM classifier described in Section 3.1. The predicted class was then recorded for each residue, together with whether a different register was used to achieve the CMO. Encouragingly, most of these residues could be correctly identified as part of modelling errors using both approaches (Figure 8), and only 34 residues were left undetected. Interestingly, while the calculation of the CMO proved to be an accurate approach for the identification of most residues found in register errors, there was a significant number of residues that could only be detected using the SVM classifier. Furthermore, only the SVM was able to identify any of the 28 residues originating from non-register related errors found in this dataset, having detected 12 of them. This suggests that while both approaches can successfully identify register errors, other kinds of modelling errors can only be found using the SVM, an indication that both methods complement each other.

**Figure 8.**
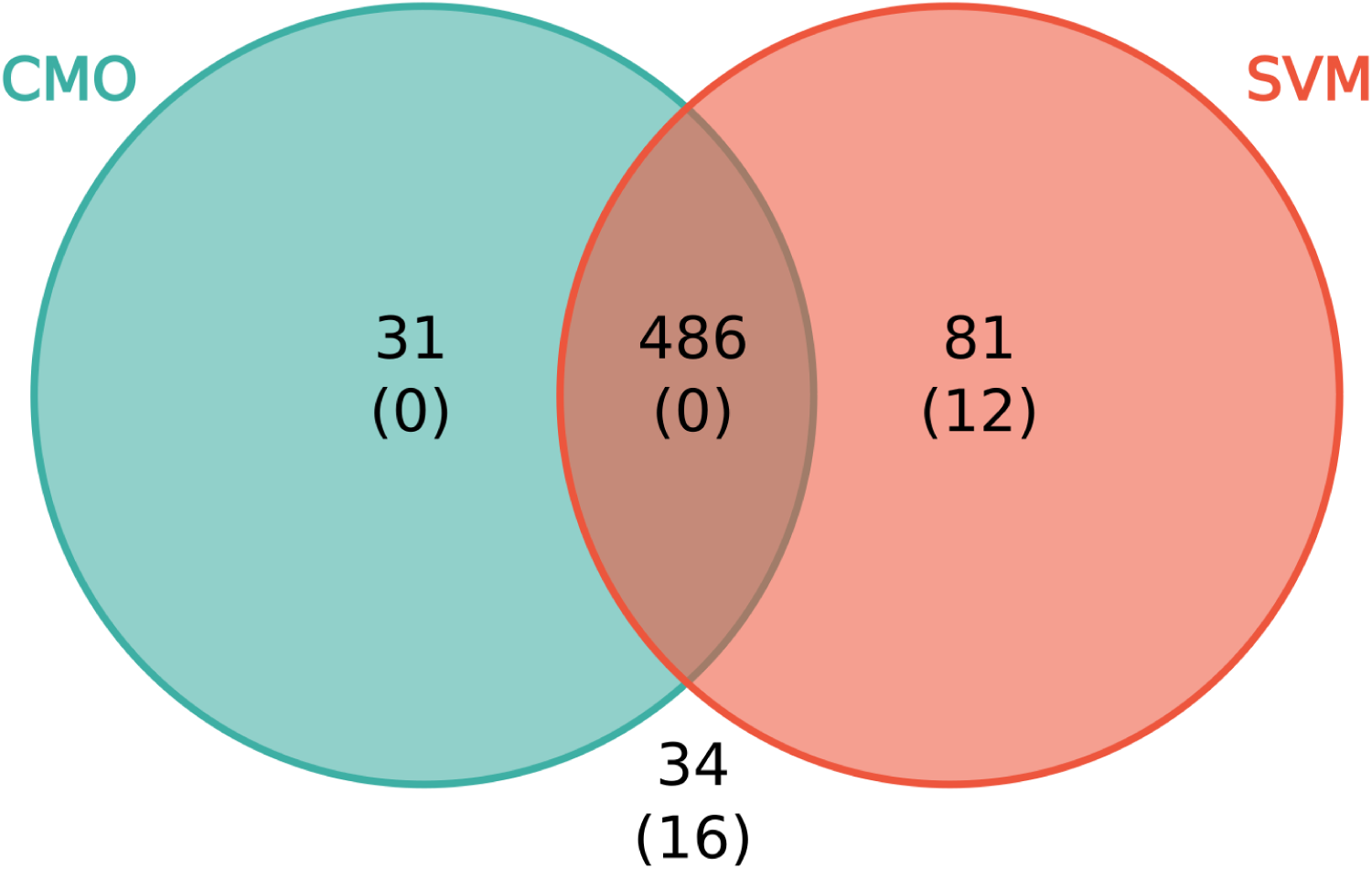
Venn diagram showing the number of residues found in modelling errors that could only be detected using the CMO approach (turquoise), the SVM classifier (orange), both methods (intersection) or neither of them (out of the circumferences). Residues found in the training set for the SVM classifier have been discarded for this analysis. The numbers of residues involved in non-register related modelling errors are shown in parentheses.

### 3.4 Identification of register errors in cryoEM structures deposited at the PDB

A new covariance-based model validation pipeline was created based on the combination of the trained SVM classifier and the contact map alignment-based sequence re-assignment methods described in previous Sections 2.4 and 2.5. For each of the residues in the input model, the pipeline outputs the classifier’s predicted probability that the residue is part of a register error and whether a different sequence register is necessary to achieve the CMO in the contact map alignment step. This pipeline was integrated into the python package for the manipulation of covariance data, conkit (Simkovic et al., 2017), in the form of a new command line option “conkit-validate”. In order to assess the performance of the proposed pipeline, a large-scale analysis was carried out with the same dataset used to analyse the performance of the validation tool checkMySequence (Chojnowski, 2022). That analysis localised possible register errors in 419 protein chains found across 246 structures deposited in the PDB with resolutions varying between 2.5Å and 4.9Å, all of them determined using cryo-EM. Given computational restrictions, and the large size of some targets, it was possible to obtain AlphaFold 2 predictions for 364 chains, representing 170 unique sequences, as described in Section 2.6. The new pipeline conkit-validate was then used to analyse all the chains in the dataset, which revealed a total of 541 possible register errors for which the CMO was achieved using an alternative sequence register, of which 230 were found to be unique when taking in consideration the fact that some sequences are represented several times in the dataset in homomeric structures. Figure 9 shows characteristics of these 230 putative register errors.

**Figure 9.**
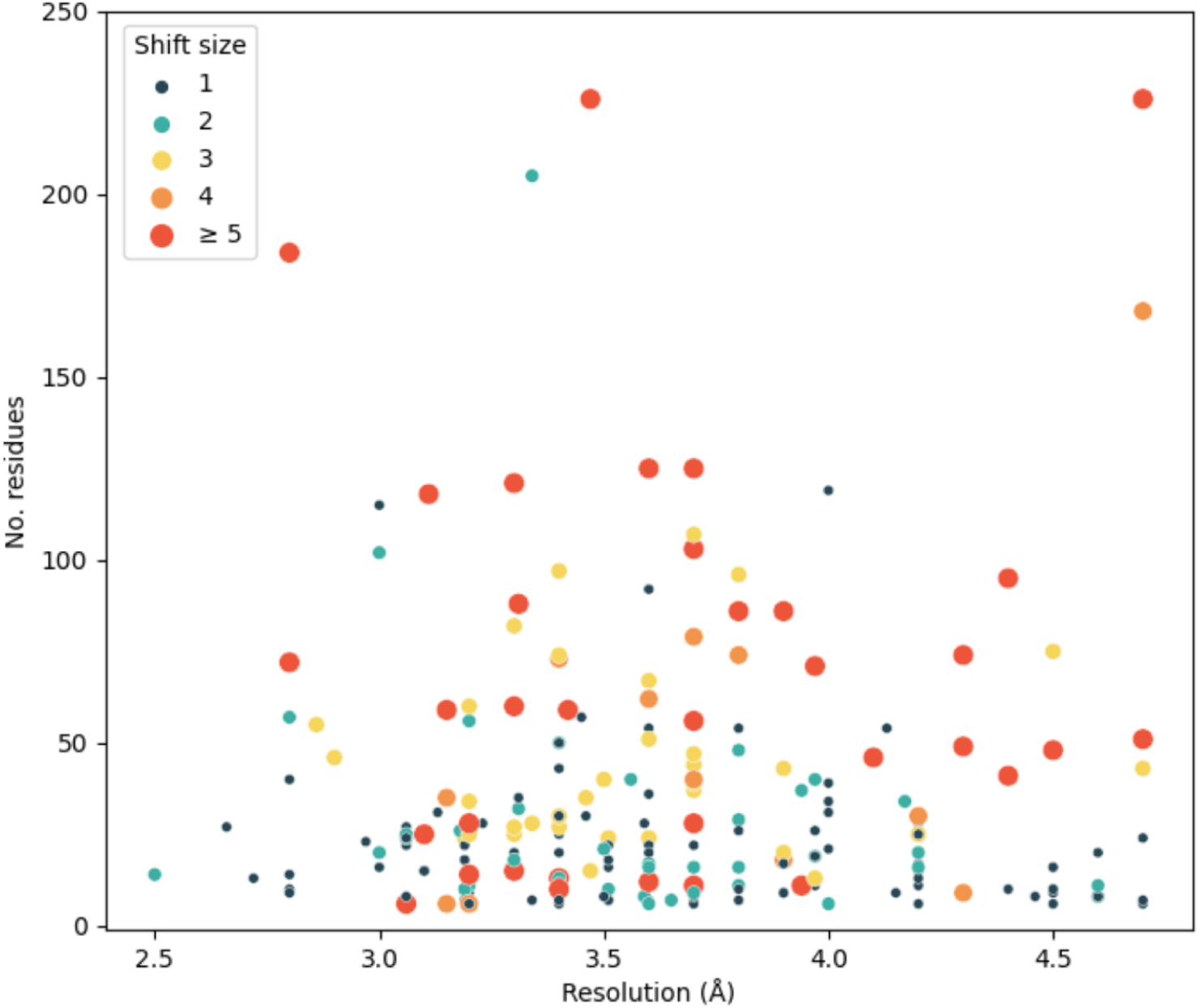
Characteristics of putative sequence register errors found in the dataset of cryoEM structures deposited in the PDB. Each point represents a predicted register error found in a chain in the dataset. Errors within 5 residues of each other were merged together into a single data point. For cases where errors are present in the same range of residues among several homomers, only one error is displayed. The vertical axis indicates the number of residues affected by the register error, which varies between 5 and 720 residues: four register errors with more than 250 residues were found, but have been omitted for clarity. The horizontal axis shows resolution, which varies between 2.5 and 4.9 Å. The colour and size of the point depict the average sequence shift observed in the error: one residue (dark blue), two residues (turquoise), three residues (yellow), four residues (light orange) and five residues or more (dark orange).

While most register errors consisted of shifts of one or two residues affecting 50 or fewer residues, a total of 18 possible register errors consisting of 100 residues or more were found across 15 different structures (Table 2). Encouragingly, further analysis of these large errors revealed, for 10 of them, the existence of at least one entry deposited in the PDB with the register that achieved the CMO. With the exception of *{PDB CODE 1}* (predicted incorrect register) and *{PDB CODE 2}* (register proposed by conkit-validate) which were deposited at 3.34 and 3.5 Å respectively, later structures with the register scoring the CMO were determined at higher resolution. Additional assessment of these models was carried out by calculating the Fourier shell correlation (FSC) between the range of residues and the density maps using REFMAC5 (Nicholls et al., 2018). Interestingly, in four out of 10 cases the structure with the alternative sequence register achieving the CMO was observed to also achieve the highest FSC of the pair. While the FSC is a well-established valuable metric for the agreement between the model and the map, it can sometimes be hard to interpret due to variations in local resolution or the effects of map sharpening. That this conventional model-to-map fit measurement does not support some of the models with the alternative register found to achieve the CMO highlights these limitations.

**Table 2.**
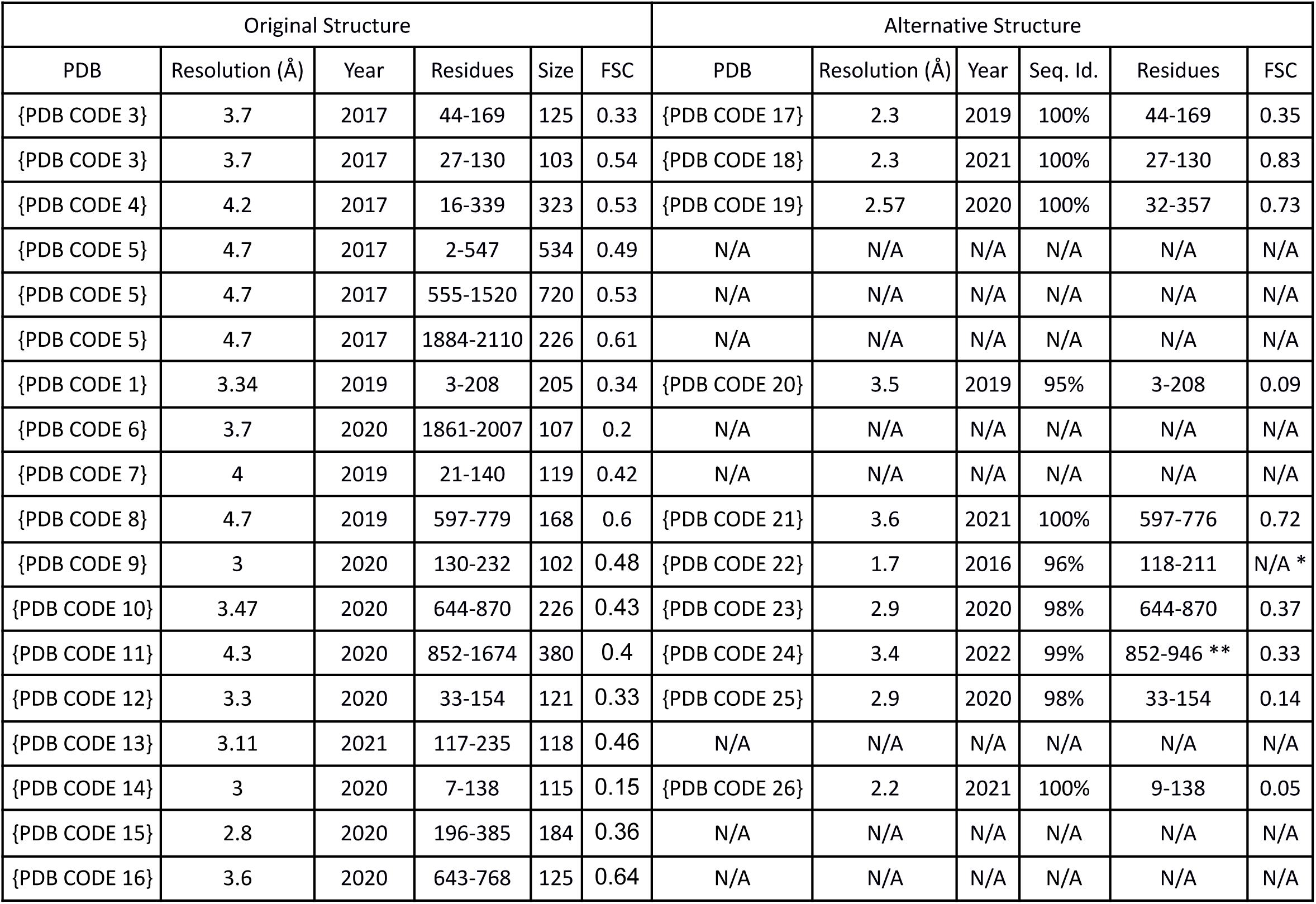
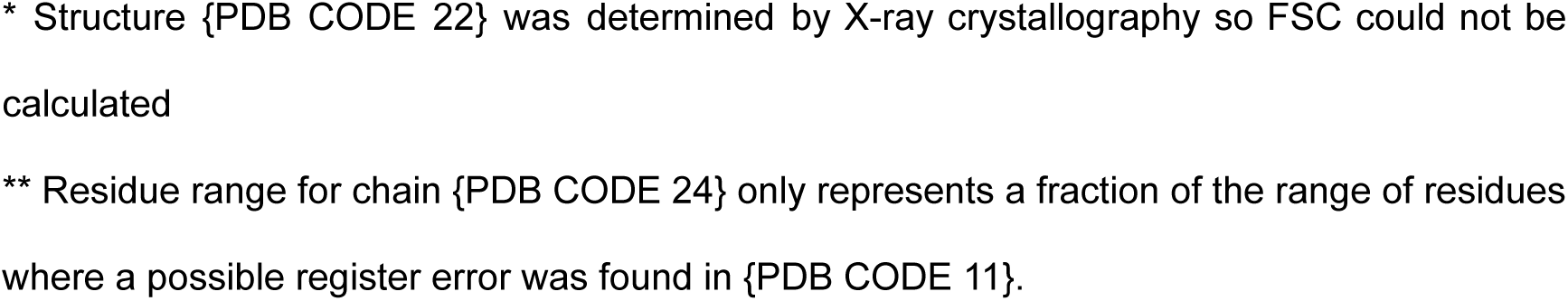
List of register errors spanning 100 residues or more that were found using conkit-validate in the checkMySequence benchmark dataset. Structures listed under “Original Structure” correspond with the structures where an error was found using conkit-validate, and those listed under “Alternative Structure” correspond with a PDB deposition for the same structure where the register matches the one found to achieve the CMO. Residue ranges and the size of the register error might differ due to the presence of missing residues. FSC refers to the fourier shell correlation between the specified range of residues and the density map. For each pair of structures, the calculation of the FSC was done using the resolution at which the structure with the lowest resolution was deposited. Sequence identity was calculated using all residues in both chains.

| Original Structure |  |  |  |  |  | Alternative Structure |  |  |  |  |  |
| --- | --- | --- | --- | --- | --- | --- | --- | --- | --- | --- | --- |
| PDB | Resolution (Å) | Year | Residues | Size | FSC | PDB | Resolution (Å) | Year | Seq. Id. | Residues | FSC |
| {PDB CODE 3} | 3.7 | 2017 | 44-169 | 125 | 0.33 | {PDB CODE 17} | 2.3 | 2019 | 100% | 44-169 | 0.35 |
| {PDB CODE 3} | 3.7 | 2017 | 27-130 | 103 | 0.54 | {PDB CODE 18} | 2.3 | 2021 | 100% | 27-130 | 0.83 |
| {PDB CODE 4} | 4.2 | 2017 | 16-339 | 323 | 0.53 | {PDB CODE 19} | 2.57 | 2020 | 100% | 32-357 | 0.73 |
| {PDB CODE 5} | 4.7 | 2017 | 2-547 | 534 | 0.49 | N/A | N/A | N/A | N/A | N/A | N/A |
| {PDB CODE 5} | 4.7 | 2017 | 555-1520 | 720 | 0.53 | N/A | N/A | N/A | N/A | N/A | N/A |
| {PDB CODE 5} | 4.7 | 2017 | 1884-2110 | 226 | 0.61 | N/A | N/A | N/A | N/A | N/A | N/A |
| {PDB CODE 1} | 3.34 | 2019 | 3-208 | 205 | 0.34 | {PDB CODE 20} | 3.5 | 2019 | 95% | 3-208 | 0.09 |
| {PDB CODE 6} | 3.7 | 2020 | 1861-2007 | 107 | 0.2 | N/A | N/A | N/A | N/A | N/A | N/A |
| {PDB CODE 7} | 4 | 2019 | 21-140 | 119 | 0.42 | N/A | N/A | N/A | N/A | N/A | N/A |
| {PDB CODE 8} | 4.7 | 2019 | 597-779 | 168 | 0.6 | {PDB CODE 21} | 3.6 | 2021 | 100% | 597-776 | 0.72 |
| {PDB CODE 9} | 3 | 2020 | 130-232 | 102 | 0.48 | {PDB CODE 22} | 1.7 | 2016 | 96% | 118-211 | N/A * |
| {PDB CODE 10} | 3.47 | 2020 | 644-870 | 226 | 0.43 | {PDB CODE 23} | 2.9 | 2020 | 98% | 644-870 | 0.37 |
| {PDB CODE 11} | 4.3 | 2020 | 852-1674 | 380 | 0.4 | {PDB CODE 24} | 3.4 | 2022 | 99% | 852-946 ** | 0.33 |
| {PDB CODE 12} | 3.3 | 2020 | 33-154 | 121 | 0.33 | {PDB CODE 25} | 2.9 | 2020 | 98% | 33-154 | 0.14 |
| {PDB CODE 13} | 3.11 | 2021 | 117-235 | 118 | 0.46 | N/A | N/A | N/A | N/A | N/A | N/A |
| {PDB CODE 14} | 3 | 2020 | 7-138 | 115 | 0.15 | {PDB CODE 26} | 2.2 | 2021 | 100% | 9-138 | 0.05 |
| {PDB CODE 15} | 2.8 | 2020 | 196-385 | 184 | 0.36 | N/A | N/A | N/A | N/A | N/A | N/A |
| {PDB CODE 16} | 3.6 | 2020 | 643-768 | 125 | 0.64 | N/A | N/A | N/A | N/A | N/A | N/A |

\* Structure {PDB CODE 22} was determined by X-ray crystallography so FSC could not be calculated
\*\* Residue range for chain {PDB CODE 24} only represents a fraction of the range of residues where a possible register error was found in {PDB CODE 11}.

Among the structures where an alternative deposition was found in the PDB with the proposed register, a 326 residue-long anti-CRISPR protein solved at 4.2Å (5XLP) was found to contain a possible register error, corresponding to the entire structure having been shifted by 10 residues towards the C-terminus. Interestingly, a structure for the same protein exhibiting the sequence register that achieved the CMO was deposited three years later at a resolution of 2.57Å (Zhang et al., 2020). Visual inspection of both models together with their respective EM maps revealed clear improvement in the match between the model and the map in the latest structure (Figure 10). The later, corrected structure was built ab initio and it seems that there was no mention of the incorrect register in the earlier structure.

**Figure 10.**
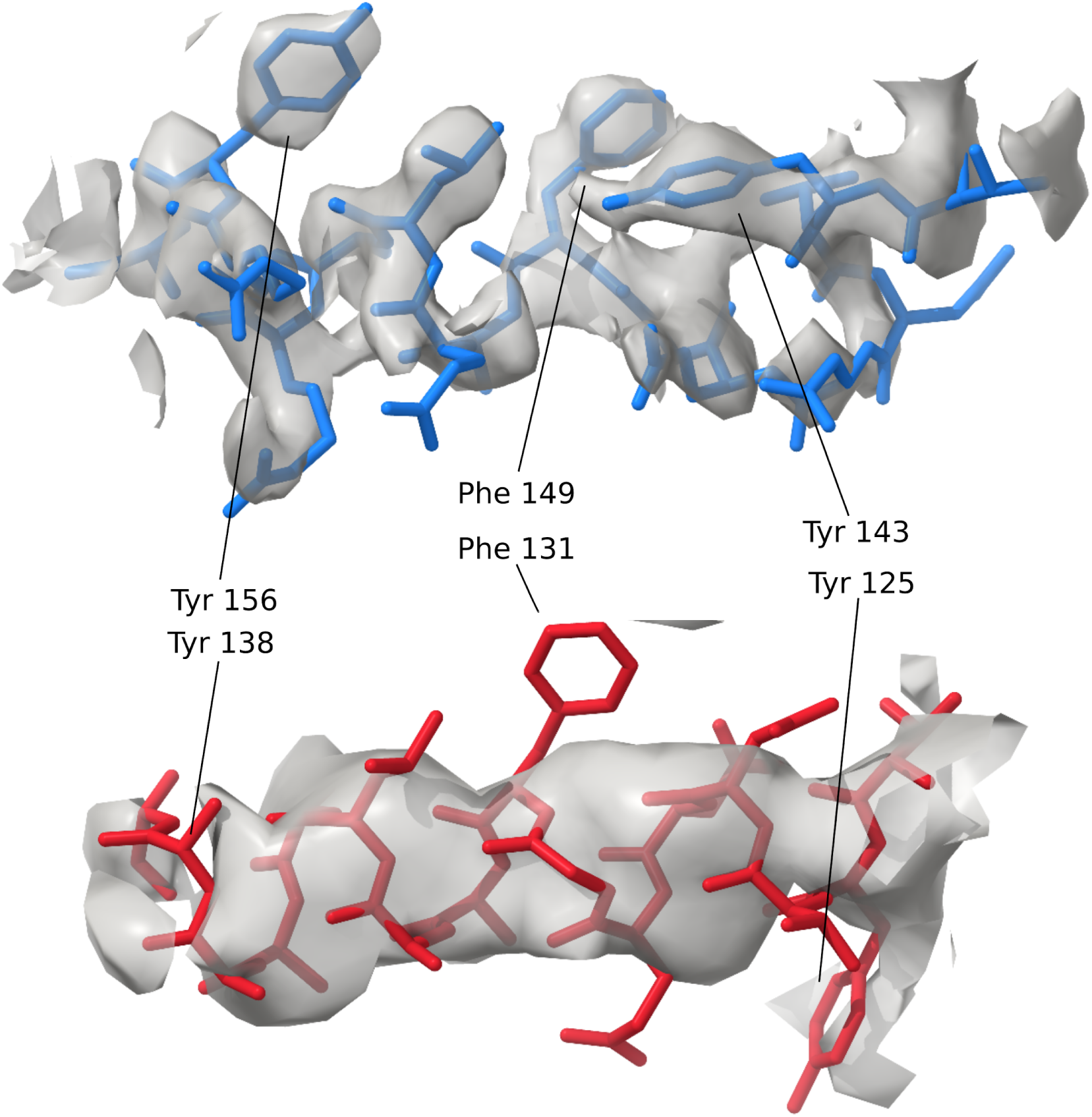
Comparison of the same section of 15 residues for a deposited structure (5XLP, chain D, resolution 4.7Å) where a possible 10-residue sequence shift was detected using conkit-validate (red model) and another deposited structure (6VQV, chain E, resolution 2.57Å) with the sequence register that achieved the CMO (blue model). The density map is represented as a transparent grey surface, and the level was set at 0.49σ for 6VQV and 0.113 for 5XLP. A mask of 3Å around both models was applied. Equivalent residues with large side chains have been highlighted for clarity (model residue numbering differs). No side chain was modelled in 5XLP for residue Tyr-138.

Another interesting case was found among the structures containing large register errors and without any similar depositions in the PDB, a human ATR-ATRIP complex (*{PDB CODE 5}*). Within this structure, three register errors each spanning more than 100 residues were predicted. While there was no reason found to disbelieve two of them, examination of the predicted error at residues 2-547 revealed a series of structural tandem repeats consisting of packed helical pairs with a cadence of approximately 40 residues. Interestingly, this matches the size of the residue shift reported by ConKit after calculating the CMO, which was of 41 residues towards the N-terminus. This suggests that the protein’s repetitive nature might have hindered the alignment of the observed and the predicted contact maps, in turn causing the false positive prediction of this unusually large register error and revealing a limitation of this approach.

### 3.5 Case study: register error found in a mycoplasma peptidase

Within the set of cryo-EM structures where a possible register error was found using conkit-validate, a sub-selection of structures was made in order to assess whether it was possible to observe evidence of these errors when inspecting the model and the density map. In order to make this assessment as unambiguous as possible, structures solved at high resolution and containing residues with aromatic side chains within the possible register error were selected. Among these structures, a mycoplasma peptidase deposited at 2.8Å was found to have a possible register error on one of its domains (*{PDB CODE 15})*. The function of this domain is not clear, although the original authors suggested that it is possibly a serine protease which function is related to the organism’s pathogenicity, specifically immune evasion.

Analysis of the validation report produced by conkit-validate (Figure 11) revealed four areas of the deposited model where the CMO was achieved using an alternative sequence register: residues 196-385, 515-524, 620-633 and 649-657. None of these errors could be verified by the existence of a different model deposited in the PDB with the register that achieved the CMO. The largest of these four errors corresponds with a 15-residue shift affecting 186 residues (residues 196-385). Interestingly, located among the residues at the end of this predicted register error, an unusual loop was found at the deposited model between residues 378-401. This loop was absent from the top ranking model produced by AlphaFold 2 (Figure 12), resulting in an otherwise structurally similar model with a different sequence register. Examination of the scores predicted by the SVM classifier for the other parts of the deposited model also revealed multiple stretches of residues predicted to be within a modelling error. While most of these stretches coincided with the portions of the model where an alternative register was suggested using the CMO approach, high scoring residues were also found in other areas of the model where the CMO did not indicate a potential register shift. Further inspection of the deposited model revealed that 25% of the residues predicted by the SVM to lie within modelling errors despite there being no issues found using the CMO approach were within 10Å of at least 5 residues for which a different register was found using the CMO. This suggests that the predicted scores for these sets of residues could have been affected by neighbouring residues found within register errors. This can occur due to the nature of the metrics used as features for the SVM classifier: inter-residue contacts and distances of a correctly modelled set of residues can still be affected by incorrectly modelled neighbouring regions.

**Figure 11.**
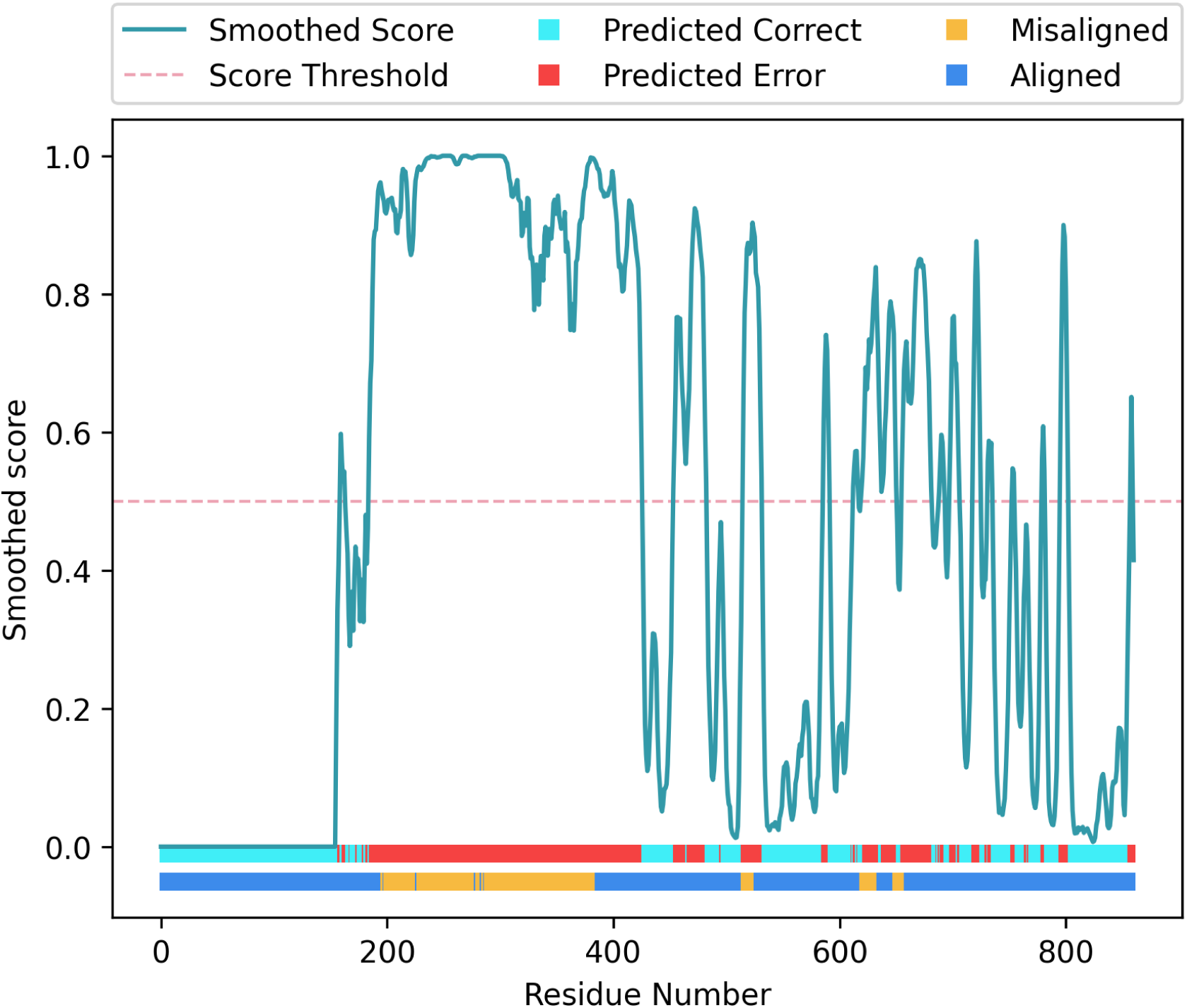
Validation report generated for structure *{PDB CODE 15}*, using conkit-validate. Scores predicted with the SVM classifier are shown as a turquoise line, and they have been smoothed using a five residue rolling average. Red dotted line shows the 0.5 score threshold. Top horizontal bar at the bottom of the figure shows for each residue position whether the predicted score was above 0.5 (red) or below (cyan). Lower horizontal bar at the bottom of the figure shows for each residue position whether the CMO was achieved using the sequence register observed in the model (dark blue) or an alternative register (yellow).

**Figure 12.**
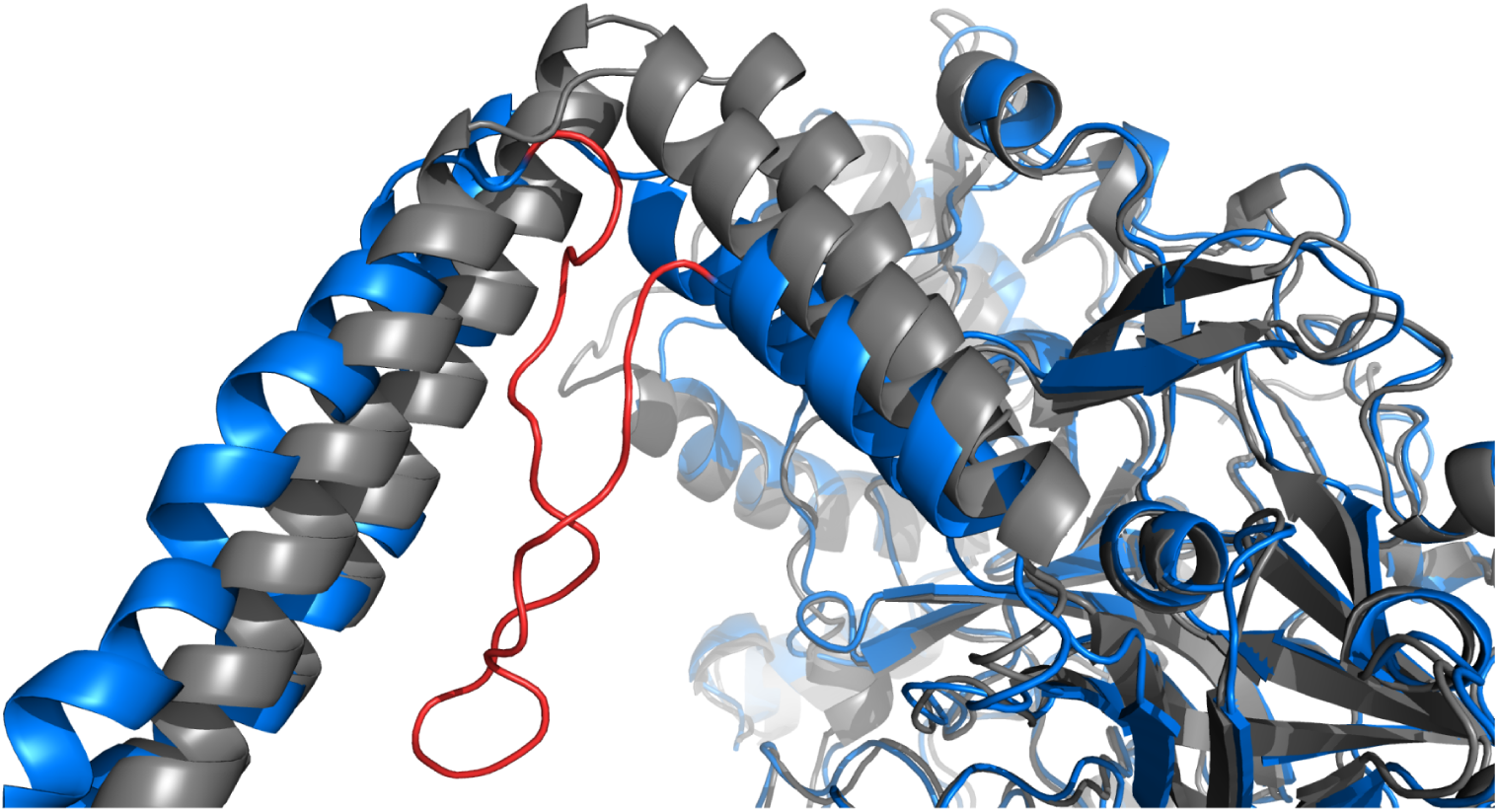
Comparison of deposited structure *{PDB CODE 15}* (blue) with the top ranked model produced by AlphaFold 2 (grey). The image focuses at the loop found in the deposited structure for the range of residues between 378 and 401 which is coloured in red.

Interestingly, analysis of the same structure using *checkMySequence* revealed a possible register error for the range of residues 613-637, which coincides with the set of residues at positions 620-633 where the CMO was achieved after shifting the sequence by one residue towards the C-terminus. Residues within this range were also assigned high predicted scores by the SVM classifier. Curiously, validation reports available for the PDB deposition showed that within this range of residues only Tyr-633 was listed as a plane outlier and a rotamer outlier. Thus, the conventional validation metrics reported by the PDB did not flag any issue with the stretch. Visual inspection of the structure and the EM map was then carried out using *Chimera-X* (Croll, 2018; Goddard et al., 2018), specifically for the range of residues where both checkMySequence and conkit-validate predicted the presence of a register error. This region of interest was then re-assigned to the new sequence register suggested by the CMO approach, using the sequence-shift tool available in *ISOLDE* (Croll, 2018). Visual inspection of these residues before and after applying this sequence shift revealed improvements in the model-map match, which is particularly evident when looking at residues with large side chains like Tyr-626 or Tyr-633 (Figure 13). Similarly, calculation of the FSC between the density map and this range of residues also revealed an improvement. After 20 cycles of jelly-body model refinement using *REFMAC5 (Nicholls et al., 2018)* on the altered structure a FSC of 0.73 was achieved, while the deposited structure registered a FSC of 0.46, an indication that in this case the conventional model-to-map fit supports the alternative register.

**Figure 13.**
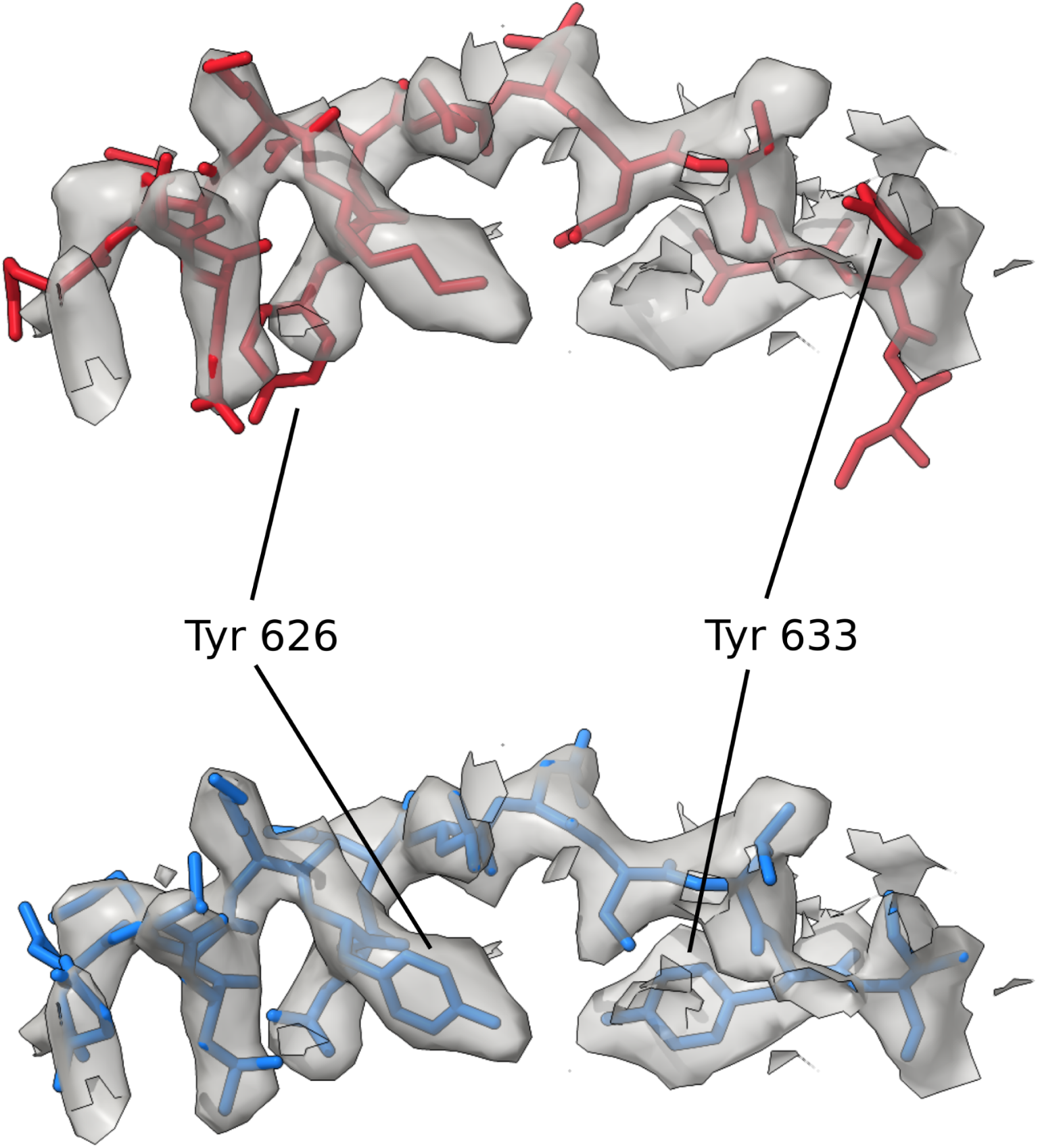
Detailed view of the section of the deposited model where a possible sequence register error was detected using conkit-validate. The density map is represented as a transparent grey surface, and the level was set at 4.8σ. A mask of 3Å around the model was applied. The original deposition has been coloured in red, and the structure with the sequence register suggested by conkit-validate in blue. Residues 626 and 633 have been highlighted for clarity. Error corresponds with PDB structure *{PDB CODE 15}*, residues 620-634.

### 3.6 Case study: register error found in an ion channel

Within the set of cryoEM structures used to benchmark conkit-validate, the structure of a ligand-gated ion channel was found to have a possible register error in a set of residues located at the receptor ligand binding domain. This structure was selected for further analysis in order to determine the presence of the detected register error as unambiguously as possible, due to the combination of high resolution (2.5Å) and residues with aromatic side chains within the possible error (*{PDB CODE 16}*).

Analysis of the validation report generated using conkit-validate for this structure (Figure 14) revealed that the sequence register for the range of residues 137-152 had to be altered in order to achieve the CMO. Additionally, this same set of residues was predicted to be part of a modelling error by the SVM classifier, a further indication of a potential sequence register error. While another 11 residues outside this range were also classified as part of modelling errors, none of these formed stretches of more than 4 consecutive residues, an indication that these are unlikely to be actual errors. Interestingly, analysis of the same structure using checkMySequence revealed a possible register error for the range of residues 136-155, in an agreement with the set of residues where a potential error was found with the other two methods. Examination of the validation report available for this PDB deposition revealed the presence of four rotamer-outliers within this range of residues: Gln-138, Gln-139, Arg-141 and Tyr-148.

**Figure 14.**
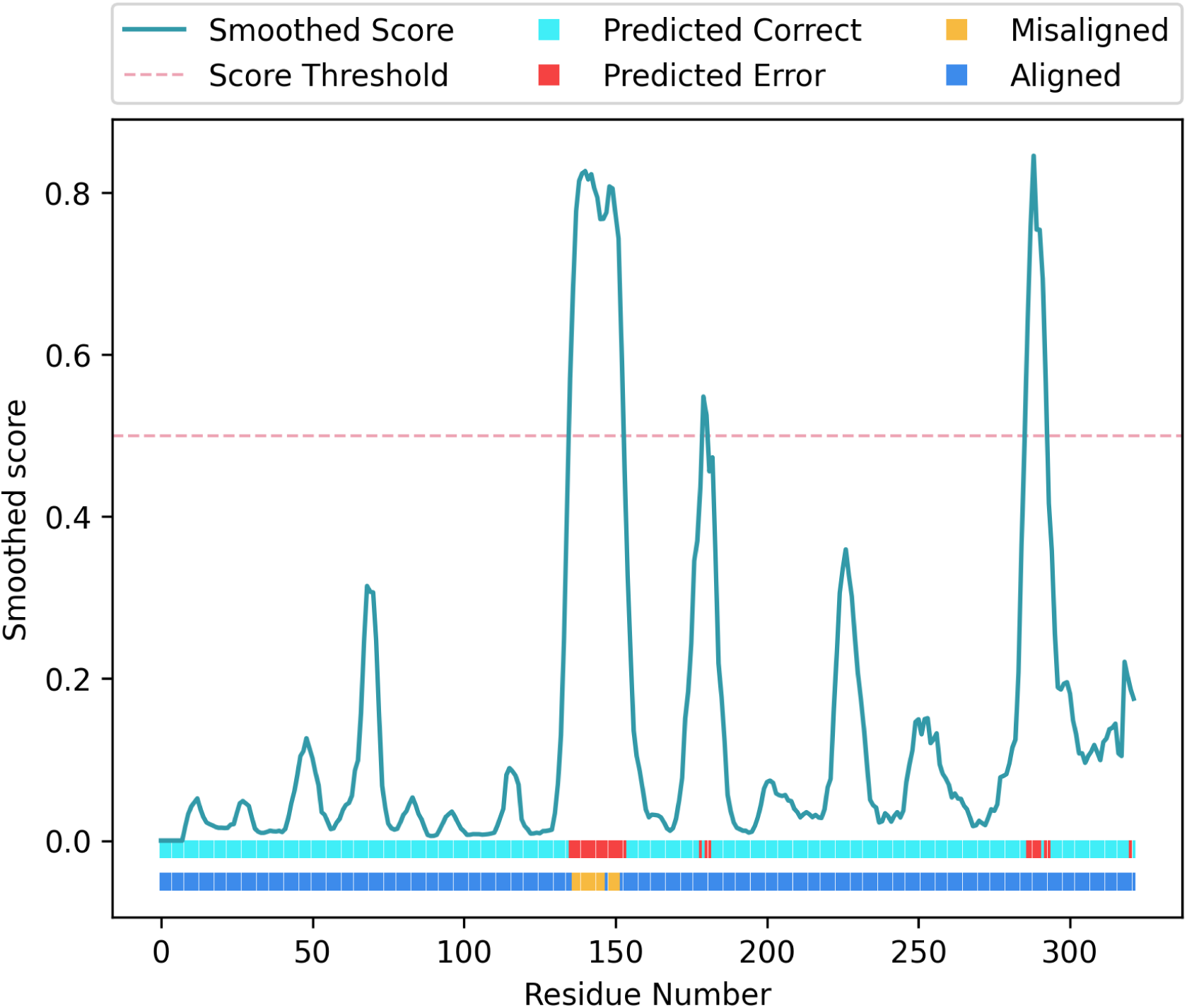
Validation report generated for structure *{PDB CODE 16}* using conkit-validate. Scores predicted with the SVM classifier are shown as a turquoise line, and they have been smoothed using a five residue rolling average. Red dotted line shows the 0.5 score threshold. Top horizontal bar at the bottom of the figure shows for each residue position whether the predicted score was above 0.5 (red) or below (cyan). Lower horizontal bar at the bottom of the figure shows for each residue position whether the CMO was achieved using the sequence register observed in the model (dark blue) or an alternative register (yellow).

The sequence register for this range of residues was then shifted by two residues towards the C-terminus using *ISOLDE* in *Chimera-X* (Croll, 2018; Goddard et al., 2018) so that it would match the register that achieved the CMO between the predicted and the observed contact maps. Visual inspection of the model before and after this modification revealed a clear improvement in the match between the side-chains of these residues and the EM map (Figure 15). After 20 cycles of jelly-body model refinement using *REFMAC5* (Nicholls et al., 2018) on the model with the alternative register, calculation of the FSC also revealed an improvement, having observed values of 0.55 and 0.76 for the deposited structure and the altered model respectively.

**Figure 15.**
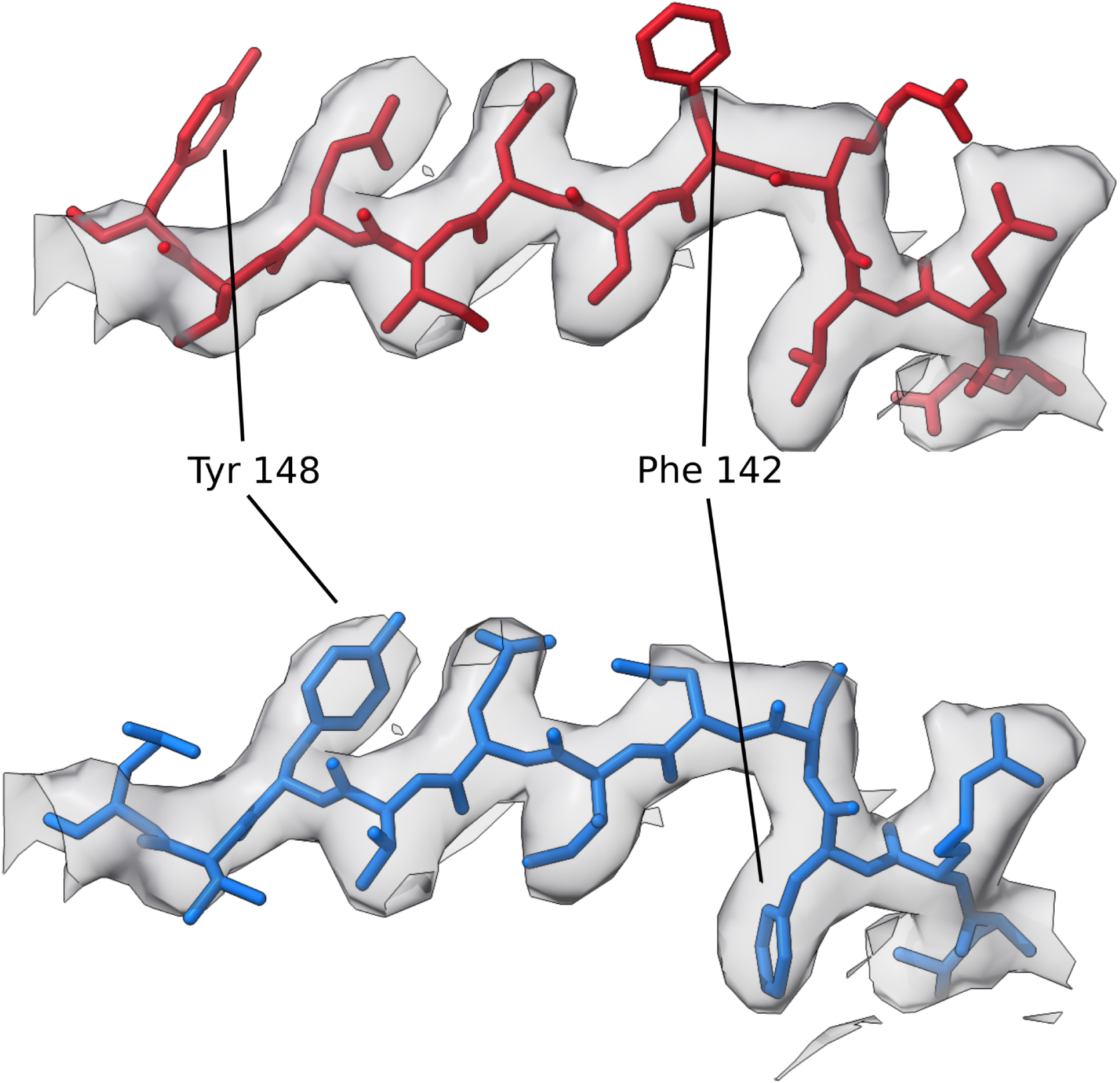
Detailed view of the section of the deposited model where a possible sequence register error was detected using conkit-validate. The density map is represented as a transparent grey surface, and the level was set to 1.35σ. A mask of 3Å around the model was applied. The original deposition has been coloured in red, and the structure with the sequence register suggested by conkit-validate in blue. Residues 148 and 142 have been highlighted for clarity. Error corresponds with PDB structure *{PDB CODE 16}* residues 138-148.

Most of the structures in this benchmarking dataset consist of structures solved using cryo-EM, deposited at a time when no previous model for the same structure had been available in the PDB. Interestingly, unlike most of the other structures in this benchmark set, this cryo-EM structure is of a ligand-gated ion channel that had already been solved by X-Ray crystallography to a resolution of 3.3Å in a previous study (*{PDB CODE 17}*) (Hilf & Dutzler, 2008). Examination of this crystal structure revealed that it shares the same sequence register with the cryo-EM structure that was found to contain a potential register error. Visual inspection of this model together with the electron density map also revealed similar features that could indicate a possible register error, particularly a poor model-map match for residues with large side-chains. A better match was then achieved after the modification of the deposited model using *ISOLDE* in *Chimera-X* to match the new sequence register proposed by conkit-validate, as revealed by further visual inspection (Figure 16). After 20 cycles of jelly-body model refinement using *REFMAC5* (Nicholls et al., 2018) on the model with the alternative register, calculation of the R_factor_ and the R_free_ also revealed an improvement, having observed values of 0.53 and 0.52 respectively, compared with scores of 0.54 and 0.54 for the deposited structure. Interestingly, an advanced search in the PDB to retrieve structures sharing at least 95% sequence identity with *{PDB CODE 16}* returned hits for 33 chains, of which 16 shared the same (predicted erroneous) sequence register for this range of residues.

**Figure 16.**
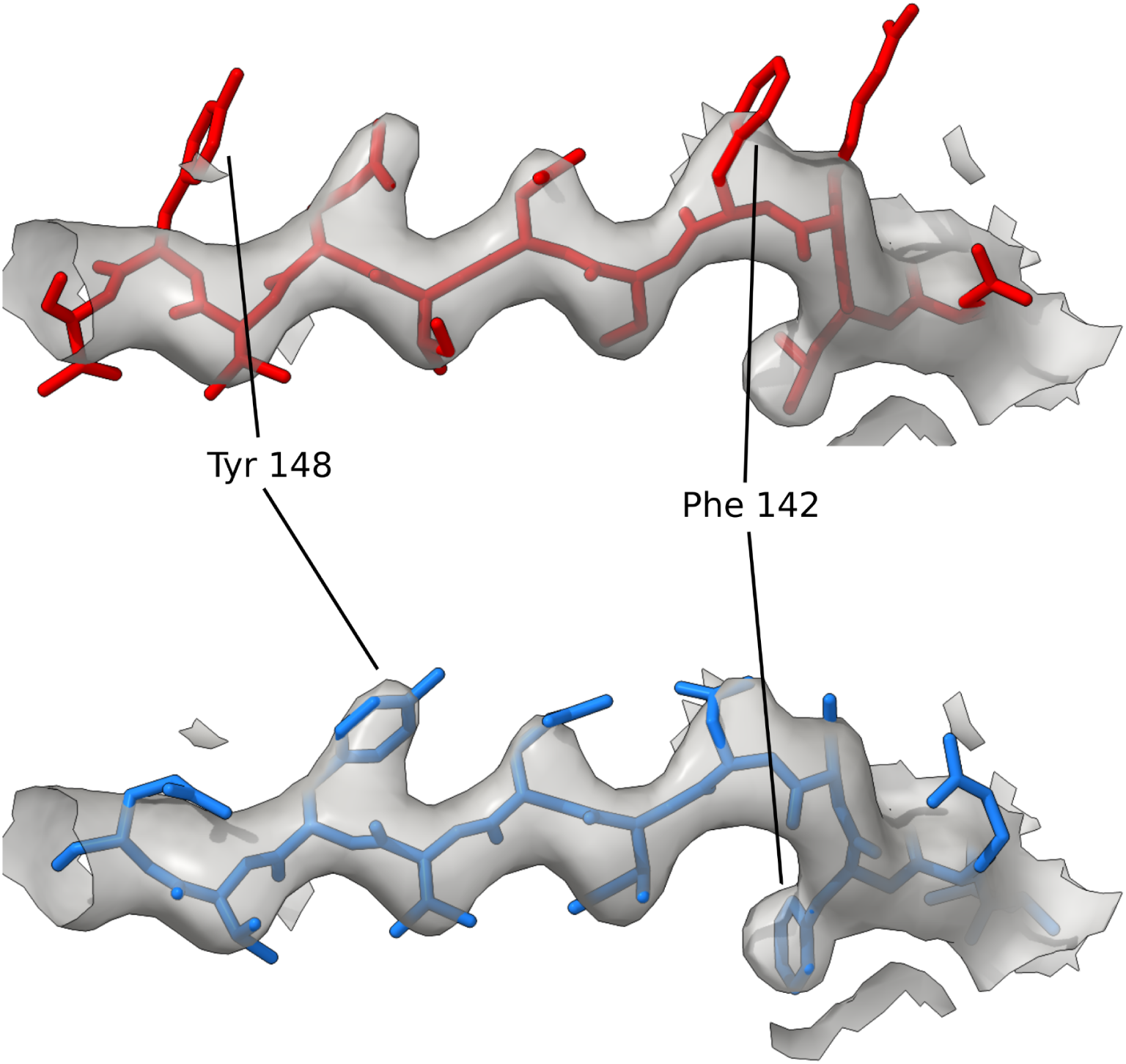
Detailed view of the section of the deposited model where a possible sequence register error was detected using conkit-validate. The density map is represented as a transparent grey surface, and the level was set to 1.06σ. A mask of 3Å around the model was applied. The original deposition has been coloured in red, and the structure with the sequence register suggested by conkit-validate in blue. Residues 148 and 142 have been highlighted for clarity. Error corresponds with PDB structure *{PDB CODE 17}* residues 139-149.

## 4. Conclusion

Here we have presented new approaches for model validation based on the use of accurate inter-residue contact and distance predictions obtained using AlphaFold 2 (Jumper et al., 2021). First, a set of new metrics were fed into a support-vector machine classifier in order to train it to detect modelling errors based on the agreement or disagreement between the observed and predicted inter-residue distances. Trained using historical data from the EM modelling challenges (Lawson et al., 2021; Lawson & Chiu, 2018), the classifier achieved an accuracy of 87% on the hold-out test set, and proved capable of detecting modelling errors among structures deposited in the PDB. Contact map alignment was then used to attempt sequence re-assignment of possible register errors: parts of the model where the contact maximum overlap was achieved using a different register to that observed in the model were marked as possible sequence register errors and the alternative register was proposed as a fix. Using this approach, it was possible to propose the correct sequence register for 87% of the register errors contained in a dataset derived from the models submitted to the EM modelling challenges. Contact map alignment has previously been used in the field of *ab initio* modelling for the selection of templates among known protein structures (Ovchinnikov et al., 2017), but to the best of our knowledge, never in the context of correcting sequence register errors.

These two new approaches were combined together into a pipeline and integrated into Conkit (Simkovic et al., 2017) as a new command line option “conkit-validate”. Future work will incorporate the method into the IRIS model validation GUI (Rochira & Agirre, 2021) that will be soon distributed with CCP4 (Winn et al., 2011) and also CCP-EM (Burnley et al., 2017) as part of the new efforts on providing tools for map and model validation. In contrast with other approaches that compare the model coordinates with the density map derived from the experimental data (Joseph et al., 2016; Liebschner et al., 2021; Pintilie et al., 2020; Ramírez-Aportela et al., 2021), our approach relies solely on the use of deep-learning based inter-residue distance predictions, which are compared with the distances observed in the model of interest. Using this new pipeline, model validation was performed for a set of cryoEM structures deposited in the PDB which were found to contain possible register errors using checkMySequence (Chojnowski, 2022). Results revealed that the use of the trained classifier in combination with the contact map alignment was successful in the detection and correction of 230 register errors which have been inadvertently deposited in the PDB. An inherent limitation of our proposed methods is the availability of high quality inter-residue distances predicted by AlphaFold2. While we don’t have reasons to believe this was the case for the set of errors shown here, we acknowledge that in cases where AlphaFold2 predictions are of poor quality - such as in regions where the predicted model scores low pLDDT - these methods may indicate potential register and modelling errors that do not really exist. In this and other respects, our method, being entirely coordinate-based, potentially has useful synergy with the map-based method of Chojnowski (2022): confident prediction of register errors can be performed in cases where these two independent methods intersect with each other. Together they represent a new generation of software that can help detect and correct the errors that even experienced structural biologists may inadvertently introduce when confronting the challenges of poorer resolution experimental data.

## References

Afonine, P. V., Klaholz, B. P., Moriarty, N. W., Poon, B. K., Sobolev, O. V., Terwilliger, T. C., Adams, P. D., & Urzhumtsev, A. (2018). New tools for the analysis and validation of cryo-EM maps and atomic models. Acta Crystallographica. Section D, Structural Biology, 74(Pt 9), 814–840.

Andonov, R., Malod-Dognin, N., & Yanev, N. (2011). Maximum contact map overlap revisited. Journal of Computational Biology: A Journal of Computational Molecular Cell Biology, 18(1), 27–41.

Baek, M., DiMaio, F., Anishchenko, I., Dauparas, J., Ovchinnikov, S., Lee, G. R., Wang, J., Cong, Q., Kinch, L. N., Schaeffer, R. D., Millán, C., Park, H., Adams, C., Glassman, C. R., DeGiovanni, A., Pereira, J. H., Rodrigues, A. V., van Dijk, A. A., Ebrecht, A. C., … Baker, D. (2021). Accurate prediction of protein structures and interactions using a three-track neural network. Science, 373(6557), 871–876.

Berman, H. M., Westbrook, J., Feng, Z., Gilliland, G., Bhat, T. N., Weissig, H., Shindyalov, I. N., & Bourne, P. E. (2000). The Protein Data Bank. Nucleic Acids Research, 28(1), 235–242.

Burnley, T., Palmer, C. M., & Winn, M. (2017). Recent developments in the CCP-EM software suite. Acta Crystallographica. Section D, Structural Biology, 73(Pt 6), 469–477.

Chiu, W., Schmid, M. F., Pintilie, G. D., & Lawson, C. L. (2021). Evolution of standardization and dissemination of cryo-EM structures and data jointly by the community, PDB, and EMDB. The Journal of Biological Chemistry, 296, 100560.

Chojnowski, G. (2022). Sequence assignment validation in cryo-EM models with checkMySequence. In bioRxiv (p. 2022.01.04.474974). https://doi.org/10.1101/2022.01.04.474974

Chojnowski, G., Simpkin, A. J., Leonardo, D. A., Seifert-Davila, W., Vivas-Ruiz, D. E., Keegan, R. M., & Rigden, D. J. (2022). findMySequence: a neural-network-based approach for identification of unknown proteins in X-ray crystallography and cryo-EM. IUCrJ, 9(Pt 1), 86–97.

Colovos, C., & Yeates, T. O. (1993). Verification of protein structures: patterns of nonbonded atomic interactions. Protein Science: A Publication of the Protein Society, 2(9), 1511–1519.

Croll, T. I. (2018). ISOLDE: a physically realistic environment for model building into low-resolution electron-density maps. Acta Crystallographica. Section D, Structural Biology, 74(Pt 6), 519–530.

Croll, T. I., Williams, C. J., Chen, V. B., Richardson, D. C., & Richardson, J. S. (2021). Improving SARS-CoV-2 structures: Peer review by early coordinate release. Biophysical Journal, 120(6), 1085–1096.

Davis, I. W., Leaver-Fay, A., Chen, V. B., Block, J. N., Kapral, G. J., Wang, X., Murray, L. W., Arendall, W. B., 3rd, Snoeyink, J., Richardson, J. S., & Richardson, D. C. (2007). MolProbity: all-atom contacts and structure validation for proteins and nucleic acids. Nucleic Acids Research, 35(Web Server issue), W375–W383.

Emsley, P., & Cowtan, K. (2004). Coot: model-building tools for molecular graphics. Acta Crystallographica. Section D, Biological Crystallography, 60(Pt 12 Pt 1), 2126–2132.

Goddard, T. D., Huang, C. C., Meng, E. C., Pettersen, E. F., Couch, G. S., Morris, J. H., & Ferrin, T. E. (2018). UCSF ChimeraX: Meeting modern challenges in visualization and analysis. Protein Science: A Publication of the Protein Society, 27(1), 14–25.

Hilf, R. J. C., & Dutzler, R. (2008). X-ray structure of a prokaryotic pentameric ligand-gated ion channel. Nature, 452(7185), 375–379.

Hooft, R. W. W., Vriend, G., Sander, C., & Abola, E. E. (1996). Errors in protein structures. Nature, 381(6580), 272–272.

Joseph, A. P., Malhotra, S., Burnley, T., Wood, C., Clare, D. K., Winn, M., & Topf, M. (2016). Refinement of atomic models in high resolution EM reconstructions using Flex-EM and local assessment. Methods, 100, 42–49.

Joseph, A. P., Olek, M., Malhotra, S., Zhang, P., Cowtan, K., Burnley, T., & Winn, M. D. (2022). Atomic model validation using the CCP-EM software suite. Acta Crystallographica. Section D, Structural Biology, 78(Pt 2), 152–161.

Jumper, J., Evans, R., Pritzel, A., Green, T., Figurnov, M., Ronneberger, O., Tunyasuvunakool, K., Bates, R., Žídek, A., Potapenko, A., Bridgland, A., Meyer, C., Kohl, S. A. A., Ballard, A. J., Cowie, A., Romera-Paredes, B., Nikolov, S., Jain, R., Adler, J., … Hassabis, D. (2021). Highly accurate protein structure prediction with AlphaFold. Nature, 596(7873), 583–589.

Kleywegt, G. J., & Jones, T. A. (1995). Where freedom is given, liberties are taken. Structure, 3(6), 535–540.

Laskowski, R. A., MacArthur, M. W., Moss, D. S., & Thornton, J. M. (1993). PROCHECK: a program to check the stereochemical quality of protein structures. In Journal of Applied Crystallography (Vol. 26, Issue 2, pp. 283–291). https://doi.org/10.1107/s0021889892009944

Lawson, C. L., & Chiu, W. (2018). Comparing cryo-EM structures. Journal of Structural Biology, 204(3), 523–526.

Lawson, C. L., Kryshtafovych, A., Adams, P. D., Afonine, P. V., Baker, M. L., Barad, B. A., Bond, P., Burnley, T., Cao, R., Cheng, J., Chojnowski, G., Cowtan, K., Dill, K. A., DiMaio, F., Farrell, D. P., Fraser, J. S., Herzik, M. A., Jr, Hoh, S. W., Hou, J., … Chiu, W. (2021). Cryo-EM model validation recommendations based on outcomes of the 2019 EMDataResource challenge. Nature Methods, 18(2), 156–164.

Liebschner, D., Afonine, P. V., Moriarty, N. W., Poon, B. K., Chen, V. B., & Adams, P. D. (2021). *CERES*: a cryo-EM re-refinement system for continuous improvement of deposited models. In Acta Crystallographica Section D Structural Biology (Vol. 77, Issue 1, pp. 48–61). https://doi.org/10.1107/s2059798320015879

Lüthy, R., Bowie, J. U., & Eisenberg, D. (1992). Assessment of protein models with three-dimensional profiles. Nature, 356(6364), 83–85.

MacArthur, M. W., Laskowski, R. A., & Thornton, J. M. (1994). Knowledge-based validation of protein structure coordinates derived by X-ray crystallography and NMR spectroscopy. In Current Opinion in Structural Biology (Vol. 4, Issue 5, pp. 731–737). https://doi.org/10.1016/s0959-440x(94)90172-4

Mariani, V., Biasini, M., Barbato, A., & Schwede, T. (2013). lDDT: a local superposition-free score for comparing protein structures and models using distance difference tests. Bioinformatics, 29(21), 2722.

Nicholls, R. A., Tykac, M., Kovalevskiy, O., & Murshudov, G. N. (2018). Current approaches for the fitting and refinement of atomic models into cryo-EM maps using CCP-EM. Acta Crystallographica. Section D, Structural Biology, 74(Pt 6), 492–505.

Ovchinnikov, S., Park, H., Varghese, N., Huang, P.-S., Pavlopoulos, G. A., Kim, D. E., Kamisetty, H., Kyrpides, N. C., & Baker, D. (2017). Protein structure determination using metagenome sequence data. Science, 355(6322), 294–298.

Pedregosa, F., Varoquaux, G., Gramfort, A., Michel, V., Thirion, B., Grisel, O., Blondel, M., Prettenhofer, P., Weiss, R., Dubourg, V., Vanderplas, J., Passos, A., Cournapeau, D., Brucher, M., Perrot, M., & Duchesnay, E. (2011). Scikit-learn: Machine Learning in Python. Journal of Machine Learning Research: JMLR, 12, 2825–2830.

Pintilie, G., Zhang, K., Su, Z., Li, S., Schmid, M. F., & Chiu, W. (2020). Measurement of atom resolvability in cryo-EM maps with Q-scores. Nature Methods, 17(3), 328–334.

Ramírez-Aportela, E., Maluenda, D., Fonseca, Y. C., Conesa, P., Marabini, R., Heymann, J. B., Carazo, J. M., & Sorzano, C. O. S. (2021). FSC-Q: a CryoEM map-to-atomic model quality validation based on the local Fourier shell correlation. Nature Communications, 12(1), 42.

Rochira, W., & Agirre, J. (2021). Iris: Interactive all-in-one graphical validation of 3D protein model iterations. Protein Science: A Publication of the Protein Society, 30(1), 93–107.

Ruiz-Serra, V., Pontes, C., Milanetti, E., Kryshtafovych, A., Lepore, R., & Valencia, A. (2021). Assessing the accuracy of contact and distance predictions in CASP14. Proteins, 89(12), 1888–1900.

Simkovic, F., Thomas, J. M. H., & Rigden, D. J. (2017). ConKit: a python interface to contact predictions. Bioinformatics, 33(14), 2209–2211.

Sippl, M. J. (1993). Recognition of errors in three-dimensional structures of proteins. Proteins, 17(4), 355–362.

Vriend, G. (1990). WHAT IF: A molecular modeling and drug design program. Journal of Molecular Graphics, 8(1), 52–56.

Weiss, M. S., Diederichs, K., Read, R. J., Panjikar, S., Van Duyne, G. D., Matera, A. G., Fischer, U., & Grimm, C. (2016). A critical examination of the recently reported crystal structures of the human SMN protein. Human Molecular Genetics, 25(21), 4717–4725.

Winn, M. D., Ballard, C. C., Cowtan, K. D., Dodson, E. J., Emsley, P., Evans, P. R., Keegan, R. M., Krissinel, E. B., Leslie, A. G. W., McCoy, A., McNicholas, S. J., Murshudov, G. N., Pannu, N. S., Potterton, E. A., Powell, H. R., Read, R. J., Vagin, A., & Wilson, K. S. (2011). Overview of the CCP4 suite and current developments. Acta Crystallographica. Section D, Biological Crystallography, 67(Pt 4), 235–242.

Zemla, A. (2003). LGA: A method for finding 3D similarities in protein structures. Nucleic Acids Research, 31(13), 3370–3374.

Zhang, K., Wang, S., Li, S., Zhu, Y., Pintilie, G. D., Mou, T.-C., Schmid, M. F., Huang, Z., & Chiu, W. (2020). Inhibition mechanisms of AcrF9, AcrF8, and AcrF6 against type I-F CRISPR-Cas complex revealed by cryo-EM. Proceedings of the National Academy of Sciences of the United States of America, 117(13), 7176–7182.

